# Decoding behavior with minimal and interpretable agent models

**DOI:** 10.64898/2026.01.20.700580

**Authors:** Giorgio Nicoletti, Antonio Celani

## Abstract

Understanding how living organisms process sensory information from their surroundings and translate it into decisions is a fundamental problem across biological scales – from biochemical signalling in single-cells to neural computations in animal brains. In this work, we address this challenge by introducing a method to reconstruct general decision processes directly from behavioral observations alone. Our approach is applicable to any biological agent and does not require prior knowledge of its internal mechanisms or its environment. Our agent model is defined by a recurrent dynamics over a discrete set of internal states which encode and process sensory information, and dictate which actions to execute. We validate our method on synthetic agents and demonstrate that we can exactly recover the agent’s behavior for non-trivial tasks. Then, we infer agent models from experimental data of rats performing evidence accumulation and of mice making decisions under uncertainty and in changing environments. In both cases, very few internal states suffice to reproduce the observed behavior with high accuracy. Crucially, the immediate interpretability of the inferred dynamics allows to understand the computational process underlying decision making. Our results show that our approach provides a broadly applicable framework for understanding how general agents make decisions in complex environments.

---

Decisions are everywhere. Viruses [1], bacteria [2, 3], single-cell eukaryotes [4–6], rodents, and mammals [7, 8] must continuously make decisions to survive and thrive. Yet, understanding how biological and living systems collect, process, and translate information into actions, whether biochemically or through neural processes, remains a fundamental challenge across fields. A wide range of modelling frameworks to capture biological behavior has been proposed, from Bayesian models to artificial neural networks. However, these approaches typically make strong assumptions about the decision-making process at hand or sacrifice interpretability for predictive accuracy. In this work, we develop a general framework that allows us to infer interpretable decision-making strategies directly from behavioral observations across diverse agents and tasks.

The theory of decision-making processes provides a general mathematical language to describe behavior as the repeated interaction between an *agent* and the *environment* that surrounds it. The boundary between agent and environment is the sensorimotor interface, where information is received and decisions are executed (see Fig. 1). Now, suppose we have at our disposal a dataset of sequences of interactions between an agent and the environment. How can we infer from the data a “good” model of the decision process of the agent? In principle, an agent with a perfect memory could make decisions based upon the entire history of past interactions with the environment. In practice, living systems are constrained by finite computational resources. Thus, biological agents must compress such histories in their *internal states*, whose role is to encapsulate all the information that is available to the agent and that is relevant for the task at hand. The biological realization of the internal state can in general be a complex and very high-dimensional object, like the biochemical state of a cell [9], the activity patterns of neurons in a brain [10–12] or, in a more abstract setting, the state of an artificial recurrent neural network [13, 14].

**FIG. 1.**
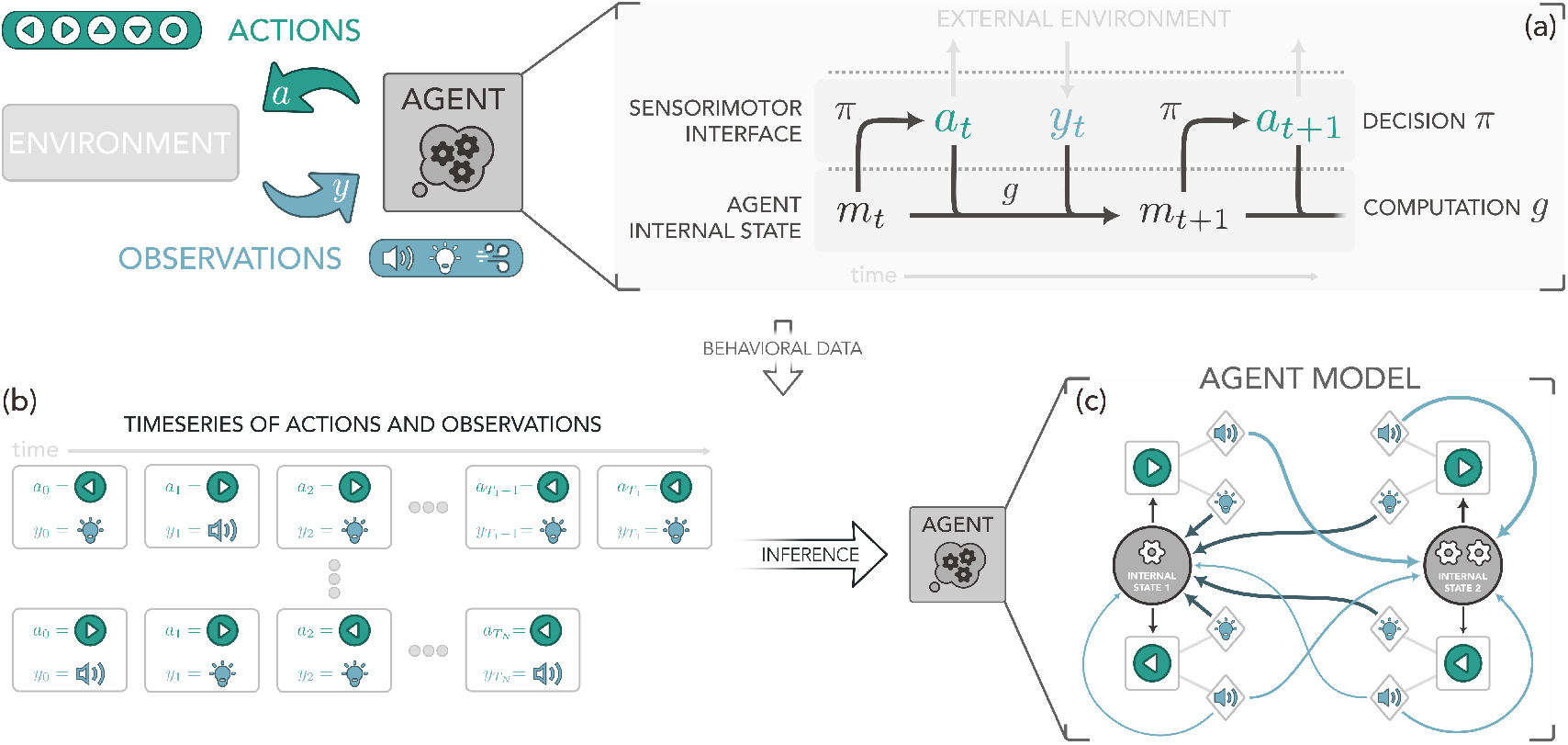
Inference of decision-making processes from behavioral trajectories. (a) We consider a generic agent that interacts with an environment via a sensorimotor interface, by taking elementary actions *a* actions (e.g., moving in space or selecting a choice) and receives observations *y* in response (e.g., sensory cues and rewards). The agent’s behavior is driven by an agent state *m*, which determines its actions through a decision process described by a probability distribution *π* over the action space. At each step of the decision process, the internal state is updated to account for new observations through an internal computation process modeled by a probability distribution *g* over the space of internal states. (b) We assume that we only have access to a series of behavioral timeseries of the agent, corresponding to sequences of action and observation pairs (*a*_*t*_, *y*_*t*_). (c) Our goal is to infer the internal model with which the agent makes decisions as a finite state controller (FSC), where the agent’s state is replaced by a finite set of discrete internal states. Its decision process is represented by stochastic policy *π*(*a*_*t*_ | *m*_*t*_) determining the action *a*_*t*_ given the current internal state *m*_*t*_, and an internal computation model *g*(*m*_*t*+1_|*m*_*t*_, *a*_*t*_, *y*_*t*_) with which the agent process information by updating its internal state depending on the previous state, the action, and the observation. In the diagram, circular nodes represent internal states, square nodes actions, and diamond nodes observations. The thickness of the arrows between nodes is proportional to the corresponding transition probability, both for the agent policy *π*(·|*m*) (gray arrows) and the computation *g*(·|*m, a, y*) (colored arrows).

Here, we employ *Finite State Controllers* (FSCs) as minimal agent models defined by a discrete set of internal states [15–19]. Formally, FSCs represent a conceptual extension of input-output Hidden Markov Models and variants alike [20–23] with the key difference that, in an FSC, actions are naturally fed back into internal transitions, allowing internal states to maintain a memory of past decisions and use it to inform future ones. The main advantage of FSCs is twofold. First, the discrete structure of internal states can be interpreted in terms of behavioral modes, governed by internal processes whose dynamics are essentially low-dimensional. This is akin to the fact that the computations of RNNs [24–27] and networks of biological neurons [28–33] can be understood in terms of transitions between discrete metastable attractors, or even belief-like representations [34–37]. Describing an agent by an FSC bypasses the need for high-dimensional fast neural dynamics by directly expressing the decision-making process through a highly coarse-grained representation of the slower processes. Furthermore, recent efforts have shown that small RNNs are able to represent a variety of cognitive tasks, suggesting that compact models of behavioral strategies can achieve the performance of more complex models [38]. Second, if the focus is on *decoding* the decision process – i.e., understanding which are the computational processes made by an agent – FSCs yield interpretable and tractable descriptions of decision strategies, making the computational processes at play explicit. These considerations lead us to ask: could an FSC with a relatively small number of internal states be a good model for the behavior of an agent engaged in a real task?

To answer this question, we develop a method to infer FSC models from sequences of interactions between the agent and the environment alone. No prior information about the environment is required, nor does the inference ever attempt to model how observations are produced – the focus is entirely on the agent. We first test our method against synthetic agents performing a variety of tasks, from arithmetic computations to spatial navigation, and show that we can efficiently discover FSCs that perfectly reproduce behavior even with a limited number of behavioral trajectories. Then, we exploit our method to infer FSCs from experimental data in rats performing an evidence accumulation task and in mice making decisions in uncertain and changing environments.

We find that inferred FSCs with few internal states not only reproduce observed behaviors with high accuracy, but also uncover the underlying computational processes driving behavior. Remarkably, in some cases, the inferred FSCs can be explicitly mapped into known decision architectures, such as exponentially smoothed counting or leaky competing accumulator models. This demonstrates that our approach provides interpretable descriptions that directly connect behavioral observations to potential neural implementations and generate testable hypotheses.

## RESULTS

### FSCs as minimal agent models

Let us describe the agent’s decisions as *actions a* ∈ 𝒜 – where *a* could be a movement in a given direction, an interaction with an object, or a measurement of some quantity – and *observations y* ∈ 𝒴 – for instance, sensory cues of different types. In general, the action *a*_*t*_ taken at time-step *t* by the agent may depend on the full history of past interactions with the environment, (*a*_0:*t*−1_, *y*_0:*t*−1_) = (*a*_0_, *y*_0_, …, *a*_*t*−1_, *y*_*t*−1_), through some probability distribution *P* ^agent^(*a*_*t*_|*a*_0:*t*−1_, *y*_0:*t*−1_).

Rather than keeping a record of past experiences, a realistic agent encodes the history in its internal state *m*_*t*_ ∈ ℳ. The agent’s behavior can then be described in terms of the probability *π*(*a*_*t*_|*m*_*t*_) of taking an action *a*_*t*_ given its current internal state *m*_*t*_, and of the recurrent computation process *g*(*m*_*t*+1_|*m*_*t*_, *a*_*t*_, *y*_*t*_), which updates the internal state by processing the information that has just been acquired. Both the *policy π* and the *internal state update g* are conditional probability distributions, so that in general the process is stochastic. The description of the decision process is completed by a choice of the initial distribution of internal states *ρ*(*m*_0_). A *model of the agent* is thus defined as the tuple ( 𝒜,ℳ, 𝒴, *ρ, π, g*), as we summarize in Figure 1a.

An FSC is an agent characterized by a discrete set of internal states |ℳ| = *M*. For the tasks considered in this work, 𝒜 and 𝒴 are also discrete sets of cardinality |𝒜| = *A* and |𝒴| = *Y*. We stress here that an FSC implicitly encodes both past observations and actions in *m*_*t*_ through the recurrency of the internal computation *g*. In this sense, it generalizes the notion of belief about the state of the external environment that characterizes Bayesian agents [39]. Further, there is an exact mapping between FSCs and linearized gated recurrent units, a gating mechanism used in recurrent neural networks (RNNs) [40, 41] (see Supplementary Information). This correspondence provides a bridge between the abstract representation of an FSC and the more biologically interpretable variables of an RNN, and generalizes input-output Hidden Markov Models [20–23] to allow internal states to retain information about past actions and use it to guide future decisions. Finally, we note that FSCs can always be made expressive enough to reproduce the behavior of an arbitrary agent over a time span *T*, albeit at the punishing price of an exponentially large number of internal states *M* ~ (*A × Y*)_*t*_. In this work, however, we focus on FSCs with relatively small *M*, following the idea that internal states should represent how the agent compresses available information, rather than performing a complex act of mimicry.

### Inference of FSCs from behavioral trajectories

We assume that we have *k* = 1, …, *N* behavioral trajectories, i.e. histories of interactions between agent and environment, each described by a sequence of actions and observations 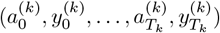 (see Figure 1b). Our goal is to infer a finite state controller that, at a given observation sequence, will reproduce the actions as closely as possible (Figure 1c). To this end, we seek FSC parameters that maximize the log-likelihood of the FSC actions given the observations, or equivalently solve the optimization problem

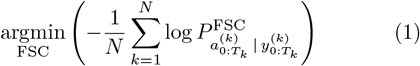

where

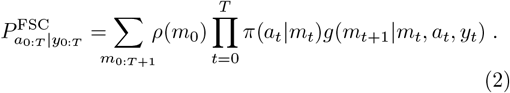

Above, the policy *π*, the internal computation *g*, and the initial distribution of the internal states *ρ*, which fully characterize the agent, are parametrized by softmax functions (see Materials and Methods).

The maximization of the likelihood in (1) is, in general, a non-convex optimization problem. Thus, in order to infer the parameters that define the FSC, we developed a global optimization method, MAPSO, whose core idea is that multiple FSCs are initialized with random parameters and then evolved according to an interacting swarm dynamics. MAPSO is based on Adaptive Particle Swarm Optimization [42], where particles are FSCs, but the topology of the interactions changes with the metric distance during the optimization. After a fixed number of iterations, the FSC that achieved the largest log-likelihood at any point during training is retained (see Materials and Methods and the SI Appendix for more details). We remark here that we do not add any regularization terms to the likelihood, such as sparsity constraints, although this is possible in principle.

Finally, we note that optimization can also be performed by a local search algorithm, e.g., gradient methods or an adapted Baum-Welch expectation maximization algorithm, augmented with random restarts. While we make no claim of computational supremacy, in our experience, MAPSO turns out to be more robust and efficient in locating global optima, at least for the cases discussed below.

### Exact inference of an arithmetic agent

We first test our inference procedure on a purely synthetic agent performing an arithmetic task. The agent can take two actions, “A” or “B”. If the agent takes A, the environment returns a random digit between 0 and 4 with equal probability. Conversely, if it takes the B action, the observation is a digit between 6 and 9. The agent has a counter that is initially set to zero. At the first step, the agent takes a random action with equal probability, observes a digit, and adds it to the counter. At all subsequent times, the agent performs a parity check on the counter and takes action A if it is even and B if it is odd. Then, it receives a new digit and adds it to the counter (Figure 2a). We want to know if our inference algorithm can retrieve the decision-making process just from a dataset of sequences of actions and observations without any information about the inner workings of the environment and of the agent.

**FIG. 2.**
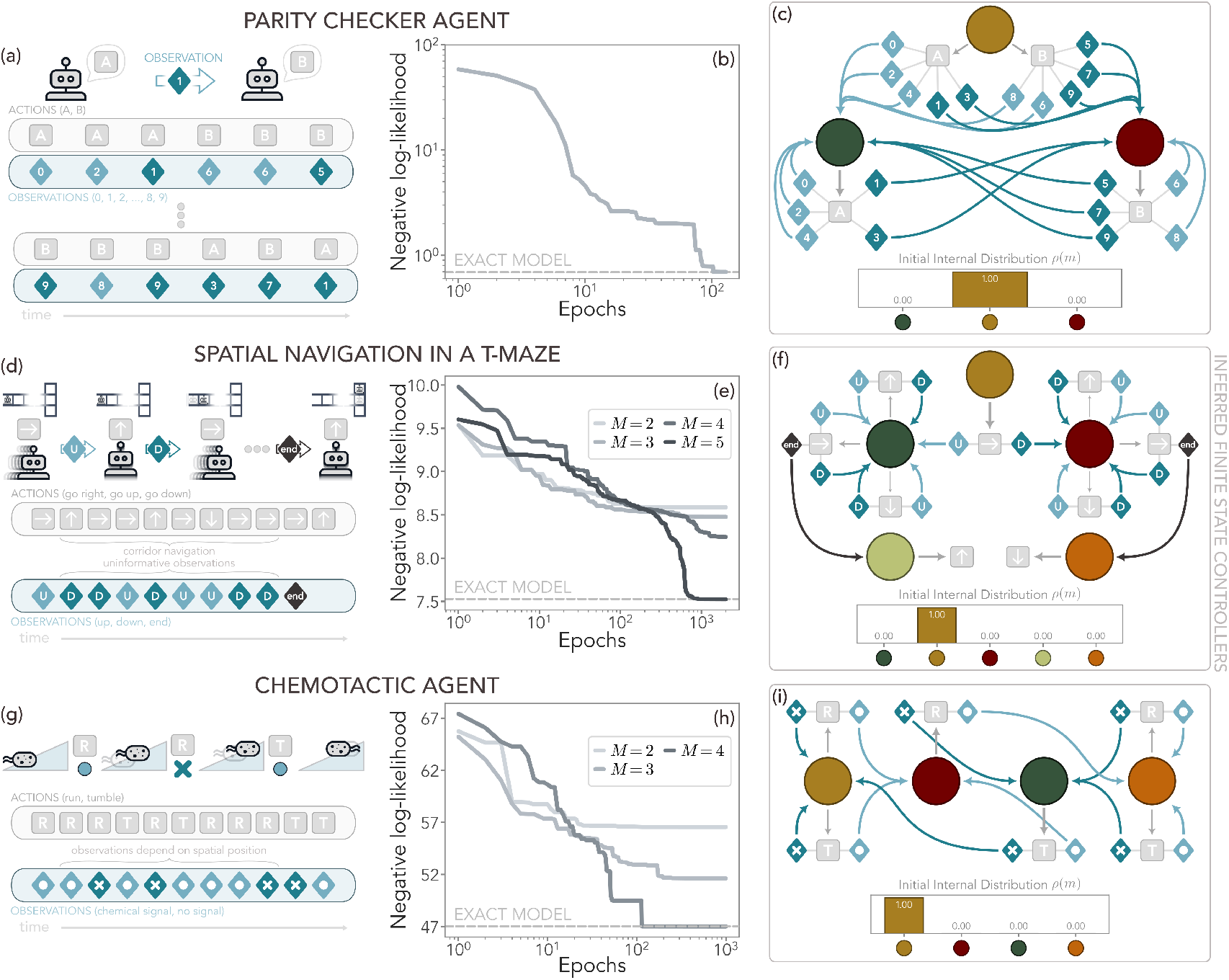
Exact inference of synthetic agents. (a) An agent can take two generic actions, “A” and “B”, and observe a digit. The agent adds the received digits to an internal counter that determines the following action according to a parity check (action A if it is even, B if it is odd). The first action is random. (b) Negative log-likelihood of an FSC with *M* = 3 internal states during training with *N* = 50 trajectories of *T* = 100 steps each. The inferred FSC perfectly reconstructs the agent’s behavior, reaching the expected negative log-likelihood (dashed gray line). (c) Inferred FSC. The 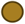 state is the initial one, where the action is taken at random 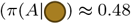, reflecting the empirical distribution). The 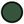 state is dedicated to the action A, whereas the 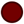 one always takes B. The transitions between the states are always deterministic and solely depend on the parity of the observation, as expected. (d) An agent performing a spatial navigation task in a T-maze 7 tiles in length. The agent enters the maze with a → action and receives an “up” (U) or “down” (D) observation with equal probability. It then either keeps moving to the right (60% probability) or steps up (↑) or down (↓) with a slight bias that depends on the first observation (30% and 10% probability, respectively). Once it reaches the T-junction, the agent chooses ↑ or ↓ solely based on the first observation, ignoring all the ones it received in the corridor. (e) Negative log-likelihood of FSCs with different *M* during training with *N* = 1000 trajectories. With *M* = 5, the inferred FSC reaches the expected negative log-likelihood of the exact model (dashed gray line). (f) The 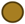 state represents the initial condition, which transitions to one of two “corridor states”, either 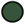 (first up observation) or 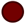 (down observation). Once it reaches the end of the corridor, the transition 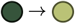 allows the FSC to correctly take ↑, while 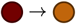 results in ↓. (g) A chemotactic agent in a one-dimensional concentration gradient. At each time, the agent may detect the presence of a chemical (observation •) or not detect it (*×*), with a probability that depends on its concentration, which in turn depends on the position in space. If in the last two steps it received the same observation, the action is chosen at random between tumble (T) which reorients the agent without any displacement, and run (R), which moves the agent in its current direction. If the last observation was• (a detection) and the previous one was *×* (no detection), then the agent runs. In the reversed case, then it tumbles. (h) Negative log-likelihood of FSCs with different *M* during training with *N* = 250 trajectories of *T* = 100 steps each. With *M* = 4, the FSC reaches the expected likelihood. (i) The internal states of the FSC encode for the four combinations of past observations: 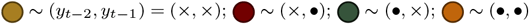 and works as a shift-register memory.

It would be natural to identify the internal states of this agent with the counter’s value, but their number would be very large if sequences are long. Are there more compact descriptions of the agent’s decision-making process? In fact, our inference algorithm discovers an exact representation of the agent’s behavior with just *M* = 3 internal states, valid for any sequence length.

In Figure 2b, we plot the negative log-likelihood during training with MAPSO from *N* = 50 behavioral trajectories of the agent of length *T* = 100. We find that the inferred FSC reaches the negative log-likelihood expected from the agent, that is the conditional entropy of the empirical distribution of actions conditioned to observations, which in this case reduces to the entropy of the initial action distribution (see Supplementary Information). In Figure 2c, we draw the inferred FSC topology. The first state, 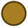 is the initial one where the action is taken at random. The other two internal states – 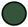 and 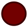 – are dedicated to taking the A and B action, respectively, and switch between them if an odd digit is observed. It is easy to check that this FSC exactly implements the agent’s decision-making rules without the need for a counter.

### Inference of a spatial navigation task

We now consider the case of a synthetic agent performing a spatial navigation task (see, e.g., [43]) that requires the exact memorization of some observations, selectively ignoring other ones, and making associations between long-past and just-received observations (Figure 2d).

The agent is stepping on a tiled T-maze and can take three actions: if it takes action →, it makes a step to the right, whereas actions ↑,↓ leave it at the same location (it bounces on the corridor walls). At the beginning of the task, the agent enters the maze from the left, always taking the → action, and receives an observation that can be either “up” or “down” with equal probability. At the subsequent steps, the agent either chooses → action most of the time ↑ or ↓ takes or with a slight bias towards the action that matches the observation received at the entrance step. As it travels along the corridor, the agent keeps receiving observations “up” or “down” with equal probability until it reaches the T-junction and receives the observation “end”. At this time, it deterministically takes ↑ if the initial observation was “up” and ↓ if it was “down”. All other observations made along the corridor do not matter – they are just noise.

In Figure 2e, we show the negative log-likelihood of the inferred FSCs with a different number *M* of internal states. With *M* = 5, the FSC perfectly reconstructs the agent’s behavior (Fig. 2f), whereas with fewer internal states the controller fails to memorize and/or make the required associations. The node 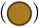 represents the “blank slate” internal state of the agent as it enters the maze. Then, the FSC branches to one of two “corridor states” depending on the initial observation, and there it remains until the junction is reached. 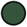 remembers the first “up” observation, so that the corresponding policy has a slight bias in the up direction. After the “end” observation, the FSC switches from 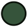 to 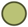, where the up action is taken deterministically. The other states, 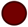 and 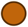, represent instead the first “down” observation, allowing the FSC to perfectly match the agent’s behavior.

### Inference of a chemotactic agent

Finally, we consider a synthetic chemotactic agent that moves in a one-dimensional concentration gradient. At each time, the agent may detect the presence of a chemical (*y*_*t*_ = •) or not (*y*_*t*_ = *×*) with a probability that depends on its position, and can either decide to run – keep on moving in its current direction, *a*_*t*_ = *R* – or tumble – randomly reorient without moving, *a*_*t*_ = *T*. The agent chooses the next action based on the last two observations according to the following policy. Assigning value *y* = 1 to the detection (*•*) and *y* = 0 to no detection (*×*), the agent computes the difference between the last two observations *y*_*t*−1_ − *y*_*t*−2_. If the difference is positive, then the agent chooses to run, if it is negative it tumbles, and finally if it is zero, it chooses at random with equal probability (see Figure 2g). At the beginning of the task, we assume that the agent has not detected a chemical in the last two time-steps. We remark that for this agent the histories of actions and observations are fully entangled: the probability of receiving an observation depends on the position, which in turn depends on the entire past history of actions, each of which depends on previous observations, and so on and so forth.

In Figure 2h, we show that the negative log-likelihood of the inferred FSCs for *M* = 4 reaches the value expected for the chosen agent. The corresponding FSC is shown in Figure 2g. Inspecting the structure of the transitions between internal states, one realizes that the nodes exactly encode for the ordered pair of last observations and the FSC works as a shift-register memory: new observations enter to the right of the register while older observations are pushed to the left and eventually out of it.

We note in passing that if the internal computation matrices are low-rank, then the likelihood is invariant under some linear mixing of the internal states (which is not just a trivial permutation), leading to multiple FSC with exactly the same behavior. This lack of identifiability indeed takes place for the chemotactic agent and is discussed in detail in the Supplementary Information.

### Evidence accumulation in rats

We now consider an experimental paradigm where rats are trained to perform evidence accumulation during an auditory discrimination task (data from Ref. [44]). During each trial, rats initially insert their nose into a central point for a fixed amount of time (1.5s, signalled by a light-emitting diode). During this time, two trains of randomly timed auditory clicks are played simultaneously, both to the left and to the right of the rat, with different click rates and total durations. At the end of the fixation period, the rat decides to poke either to the left or to the right, and receives a water reward if it turns towards the side where more clicks were played (Figure 3a). The sum of the left and right click rates is fixed at 40Hz, with subsequent trials varying in difficulty depending on the ratio of the click rates.

**FIG. 3.**
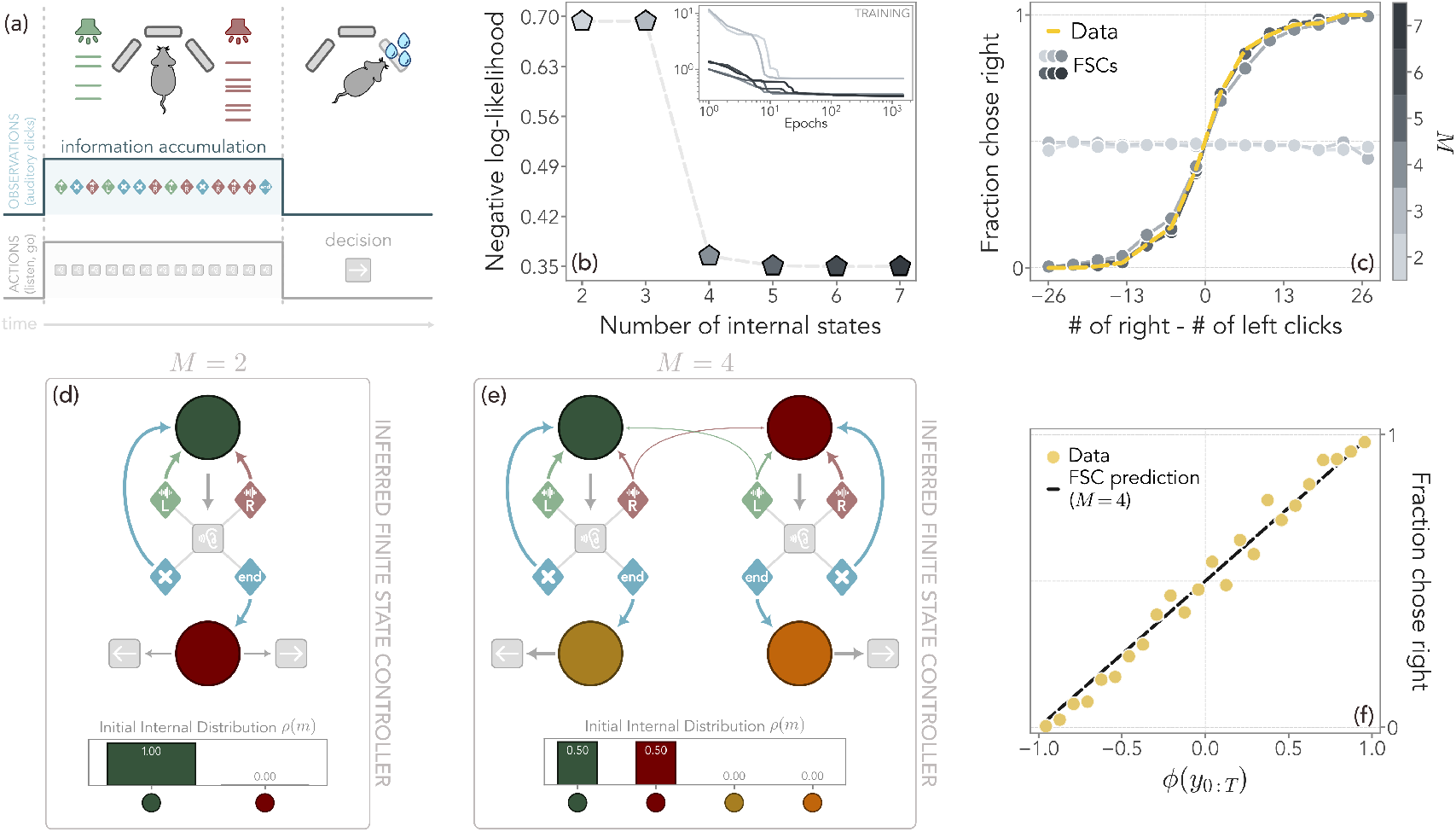
Evidence accumulation in rats during perceptual decision making. (a) Sketch of the experiment. During each trial, rats fixate their nose into a central port for 1.5s. During this time, corresponding to the action “listen”, trains of randomly timed auditory clicks are played both to the left and to the right, with different frequencies and durations (between 0.2s and 1*s*) which represent three observations: left, right, or silence (“*×*” in the figure). After nose fixation ends, the rat receives an “end” observation, and pokes in one of the side ports (actions ← or →). If it turns to the side where more clicks have played, it receives a water reward. Data of approximately 4500 trials (half for training, half for validation) from [44]. (b) Negative log-likelihood over the validation set of the inferred FSCs with different numbers of internal states. For *M* ≥ 4, a two-layer structure, which reproduces the structure of the experiments, was imposed a priori to speed up convergence. Inset: negative log-likelihood over the training set during inference. (c) At *M* ≥ 4, FSCs (grayscale dots) accurately reproduce rats’ performance (dashed sand line, averaged across trials and individuals) in choosing right as a function of the difference between the number of clicks. A further improvement can be seen for *M* = 5 onward. (d) Inferred FSC with *M* = 2 internal states. The FSC learns to take a ← or → action only after the “end” observation, but cannot perform any computation due to the lack of internal states, and thus must choose at random. (e) Inferred FSC with *M* = 4 internal states and a fixed two-layer structure. The two internal states on the top 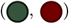 are responsible for taking the “listen” action, acting as a computation layer where the agent accumulates evidence. The two at the bottom 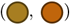 decide whether to go left or right. The state 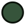 remains unchanged if a click is heard on the left or no click is heard, whereas it switches to 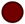 with a probability *ϵ* ≈ 0.2. If “end” is heard, the trial ends and 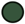 switches with probability one to 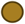, where the agent chooses to go to the left. The situation is reversed for the states 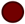 and 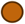, the latter of which goes to the right. (f) It can be shown that this FSC is computing an exponential smoothing of the difference between right and left observations *ϕ* with a smoothing factor *ϵ* (Eq. (3). Indeed, the fraction of times that the rats choose to go right is well-predicted by its value at the end of the trial. Since *ϕ* is captured by the internal state 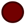, the FSC prediction simply corresponds to the bisector of the plane.

From the behavioral trajectories, we can discretize time (we set Δ*t* = 1*/*40s) and define four different observations and three actions. At each timestep, as long as light is on, the agent may only take the action “listen” and receive a “left” (*y* = − 1) or “right” (*y* = 1) observation if it hears a click on the corresponding side, or a “silence” observation (*y* = 0) if it hears neither. When the light-emitting diode turns off, the agent receives an “end” observation and can take one of the two actions “go left” or “go right”.

Among the data in [44], we extracted the behavioral data of three rats, each across two different days. To average out noise in the rats’ behavior and accurately sample different task difficulties, we retained approximately 4500 trials, with 50% used for inferring the FSC parameters and 50% to validate their performance. In Figure 3b, we show the negative log-likelihood of the best-performing FSCs for different numbers of internal states *M*. Remarkably, we find a sharp decrease at *M* = 4, after which the performance quickly plateaus. Indeed, the FSCs with *M* ≥ 4 closely reproduce the observed rats’ performance (Figure 3c) with slight deviations visible only for *M* = 4. This signals that an FSC with a relatively small number of internal states is expressive enough to reproduce the behavioral trajectories.

#### A small FSC recapitulates the essential features of decision-making

With *M* = 2 and *M* = 3, the inferred FSCs are trivial, as their limited internal structure can only reproduce the fixation period during which rats listen until the “end” cue (see Fig. 3d and the Supplementary Information). For example, with *M* = 2, no computation can be done in the first internal state because “left” and “right” observations always circle back to this state. When “end” is observed, the FSC transitions to the other state and can only choose “go right” or “go left” at random, regardless of the past history. This model can only reproduce the average final decision across all data, and therefore cannot accumulate past evidence. Inferred FSCs with *M* = 3 nodes similarly fall short.

The structure of the interactions for this task hints at a layered organization of the internal states: a computation layer where the agent can only take the “listen” action, and a decision layer responsible for taking the “go” actions and accessible only upon receiving the “end” signal. For computation to be possible, at least two nodes in the upper layer are required, whereas two are both necessary and sufficient for the final decision. We therefore focused on FSCs architectures with *M* − 2 computation layer nodes and 2 decision layer nodes. A posteriori, we then check that these structures are indeed those that minimize the negative log-likelihood.

Let us first focus on the simplest case, *M* = 4, shown in Figure 3e. We reiterate that, with the layered structure induced by the random interruption, only the two internal states of the top layer are used for internal computation. Despite the small number of states, this FSC is able to achieve a high accuracy in reconstructing behavior, with the 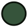 state encoding for left clicks, and 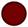 for right ones. The simplicity of the model allows to compute analytically the predicted probability of choosing an action at the end of the trial, *ϕ*(*T* + 1) = *p*(*A*_*T* +1_ = → |*y*_0:*T*_) − *p*(*A*_*T* +1_ = ← |*y*_0:*T*_), which obeys the recursion relation

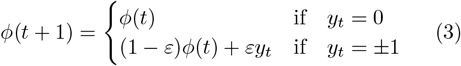

with *ϕ*(0) = 0. Note that *ϕ*(*t*) is an exponential smoothing of the difference between right and left observations, with smoothing factor *ϵ*. Here, *ϵ* ≈ 0.21 is the probability of switching between the two upper memories (see Supplementary Information). In Figure 3f, we show that indeed the fraction of trials in which the rats choose to go right is well predicted by the value of *ϕ* at the end of the trial. Thus, the simplest nontrivial FSC with *M* = 4 interprets the decision-making process as the following algorithmic procedure: compute the exponentially recency-weighted difference of right and left clicks and, at the end of the trial, bias the choice proportionally to this quantity.

#### Additional internal states allow longer memory of past events

We now seek to understand how the FSC with *M* = 5 improves upon the previous one, allowing it to reproduce the rats’ behavior with greater accuracy and to saturate the likelihood (see Figure 3b-c).

In Figure 4a, we draw the inferred FSC with five internal states. Similarly to the previous case, we find two nodes in the computation layer ( 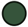 and 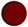) that encode the prevalence of left and right clicks, respectively. Once the trial ends, they deterministically transition to the corresponding state in the decision layer ( 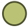 and 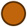) where the agent goes to the left and to the right, respectively. However, the transitions between 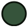 and 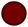 are now mediated by an intermediate node 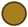. If the “end” observation is received while the FSC is in this state, it transitions to the decision layer with a slight left bias, reflecting an overall observed preference by the rats. Importantly, the 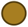 state is also the one where the FSC is initialized.

**FIG. 4.**
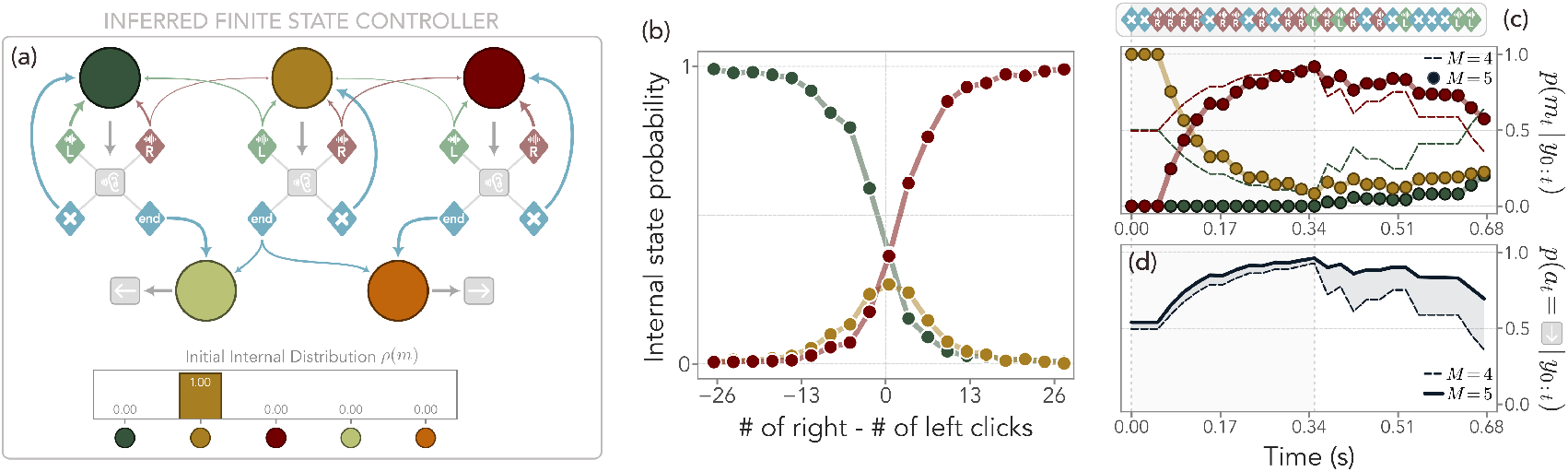
Inferred FSC with a two-layer structure and *M* = 5 internal states for evidence accumulation. (a) Similarly to the case of *M* = 4, the leftmost 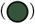 and rightmost 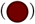 internal states in the computation layer are responsible for encoding the prevalence of left and right clicks, respectively, and go in the corresponding direction once the trial ends (respectively 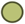 and 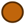). Importantly, the switch between them is regulated by the 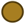 internal state, which transitions to the left or the right when the corresponding click is heard with a probability *η* ≈ 0.29. The FSC is initialized at this internal state. If the trial ends in the 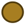 state, it transitions to either 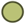 or 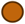 approximately at random (0.56 and 0.44, respectively). (b) Probability of occupying one of the upper internal states 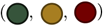 right before the trial ends, averaged over the observation sequences in the data, as a function of the difference between right and left clicks. The central state is mostly active in an uncertainty region where the difference in the number of clicks is close to zero. (c-d) For a fixed observation sequence *y*_0:*t*_, the probability of occupying a given internal state *p*(*m*_*t*_|*y*_0:*t*_) may be different between the FSC with *M* = 5 (colored dots and solid lines) and with *M* = 4 (dashed lines). In particular, when a set of congruent observations is followed by incongruent ones (e.g., a series of “right” observations first and then a series of “left” ones), the central state acts as a buffer and maintains the occupancy of the right state at higher levels. This reflects in a slower and delayed decay of the probability of taking the action “go right” if the trial were to end at time *t* (bottom panel).

In Figure 4b, we plot the probability of occupying one of the computation states 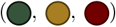 right before the trial ends as a function of the difference between the number of clicks heard on the right and on the left. When most observations are from the right, the FSC activates the 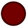 state, and vice versa for the 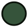 state. Crucially, the occupation of the middle 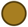 state peaks when the trial ends in a region of uncertainty, where the difference between clicks is close to zero.

This intermediate node plays an important role. In Figure 4c, we consider a fixed observation sequence *y*_0:*T*_ from the data, characterized first by a series of “right” observations followed by noisy ones where the rat mostly hears “left”. We compute the probability of occupying a given internal state at time *t* ∈ [0, *T*], 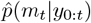, and compare it with the *M* = 4 case. We find that, after an initial transient, both FSCs reach a high level of occupancy of 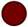, since only right clicks were heard. However, when the incongruent observations start to appear, the 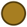 state acts as a buffer and delays the transition to 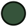, which is instead quickly occupied when *M* = 4.

We can compute the predicted probability of choosing an action at the end of the trial for this FSC as well (see Supplementary Information). Remarkably, the calculations reveal that the rats’ decision is captured by a leaky competing accumulator model (LCA) [46] with lateral inhibition. Namely, the preference for choosing right is proportional to the difference of two accumulators *ϕ*^*±*^(*t*) –favoring state 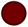 and 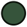, respectively – at the final time. These obey the equations

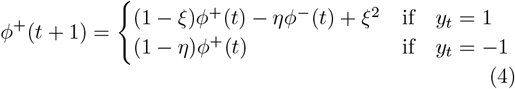

and

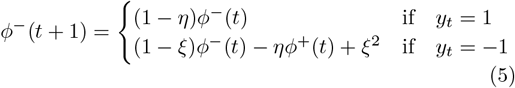

with *η* ≈ 0.27 the probability of exiting the central state 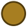, and *ξ* ≈ 0.13 the probability of exiting one of the extremal states (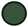 and 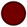). Thus, the passive accumulator – the one that contrasts with the current evidence *y*_*t*_ – inhibits the active one. Due to this lateral inhibition, the probability of taking the right action if the trial were stopped at time *t* decays more slowly, allowing the FSC with *M* = 5 to reproduce the rats’ behavior more accurately and helps identify the side on which more clicks were played, as shown in Figure 4d.

### Mice decision-making in changing environments

We now consider a different experiment, where mice are trained to make decisions in a changing and uncertain environment (data from [45]). At each step, a mouse can choose from two side ports. One of the two ports has a high probability, e.g. 0.8, of delivering a water reward (the high port), whereas the other has a probability 0.2 of rewarding the mice (the low port). After the completion of each trial, the high and the low ports have a probability 0.02 of switching. This results in blocks of consecutive trials, of variable length, where the environment is predictable, even if uncertain. When the environment switches, however, the mouse has to swiftly adapt its strategy to the change (Figure 5a).

**FIG. 5.**
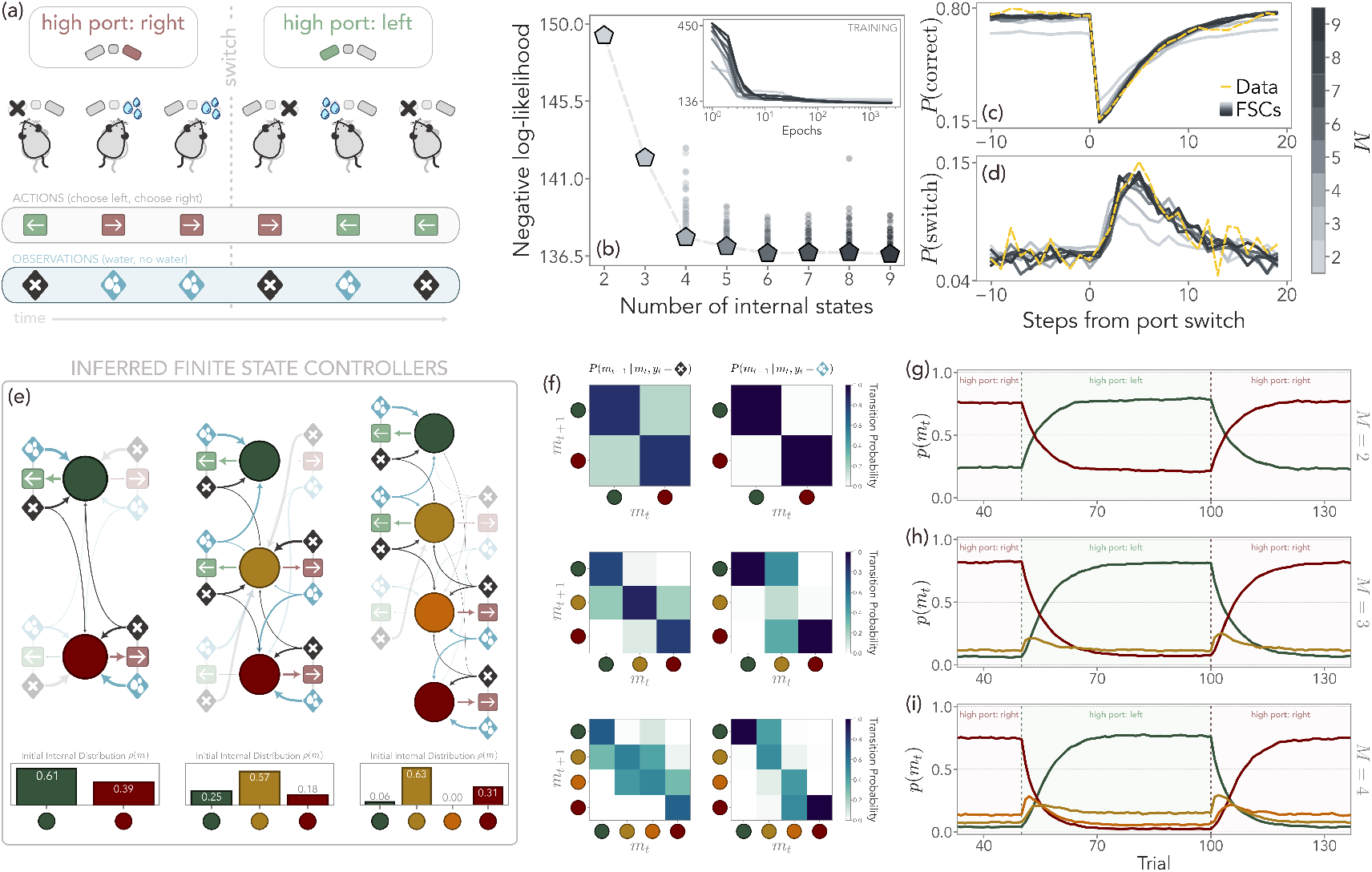
Probabilistic decision-making strategies in mice. (a) Sketch of the experiment. Mice are trained to select one of two side ports, representing two possible actions. Each port has a different probability of giving a water reward, either 80% (the high port) or 20% (the low port), corresponding to the two possible observations. At each time, the high port may randomly switch with a 2% probability, so that mice have to adjust their choices accordingly. Data of 159 behavioral trajectories (126 for training, and 33 for validation) from [45]. (b) Negative log-likelihood over the validation set of the inferred FSCs with different numbers of internal states. Pentagons represent the FSCs with minimum loss across restarts of the inference procedure, and selected restarts are represented by dots. Inset: evolution of the negative log-likelihood over the training set during inference. (c-d) The FSCs reproduce both the probability of selecting the higher reward port (*P* (correct)) and the probability of switching port selection when the high port changes (*P* (switch)), with an accuracy that increases with the number of internal states *M* and saturates from *M* = 4. (e) Inferred FSC with *M* = 2, *M* = 3, and *M* = 4. From an internal state, green arrows represent the probability of taking the action left, and red arrows the probability of choosing right. Transitions between the states depend on the observation, either when a reward is received (blue) or not (black). For ease of visualization, actions with low probability and the corresponding state transitions have a higher transparency. In all cases, we can identify two extremal states (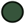 and 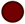) primarily encoding the left or right action, respectively. For *M >* 2, the central states ( 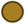 and 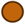) buffer the transitions between the extremal ones, similarly to Figure 4. (f) Marginalizing over the actions emphasizes the linear structure of the FSC. For *M >* 2, the FSCs have a high probability of transitioning to the central states in the absence of a reward, whereas they drift towards the extremal states when a reward is obtained. (g-i) Probability of occupying the internal states as a function of time and for a specific sequence of high ports (right, left, right), averaged over 10^4^ trajectories. With *M* = 3 and *M* = 4, the central states quickly activate when the port changes, playing the role of switch detectors. This allows the FSC to mimic the mice’s strategy and switch to the high-reward action.

From the behavioral trajectories, we can immediately identify two possible actions – “choose left” or “choose right” – and two observations – “water” and “no water”. Importantly, and differently from the previous case, the observations received by the agent are now explicitly dependent on the action it has taken. Furthermore, since any observation can follow from an action, we do not impose any a priori structure on the FSC. We consider the 159 trials reported in [45], with 80% of them used for inference and 20% for validation. In Figure 5b, we plot the negative log-likelihood of the inferred FSC for different numbers of internal states. Once more, we find that the likelihood plateaus already at relatively small values of *M*. However, at variance with the previous section, we find pronounced improvements when moving from *M* = 2 to *M* = 4.

#### Simple FSCs successfully capture the switching behavior

To quantitatively assess the ability of the FSC in capturing the decision process of the mice, we compute the probability of selecting the high reward port *P* (correct) and the probability of switching port selection when the high port changes *P* (switch), both as a function of the time from a switch [45]. We find that the inferred FSCs with *M* = 4 or above are able to reproduce the data very accurately. Remarkably, even for *M* = 2 and *M* = 3, the FSCs are still able to capture the timescales at which mice switch their behavior on average.

In Figure 5e, we draw the structure of the inferred FSCs with *M* = 2, 3, 4. In all cases, we find two extremal states (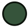 and 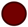) where the probability of choosing left and right, respectively, is significantly higher. This suggests that these states encode the agent’s confidence that the corresponding side is the one with a higher chance of delivering the reward. For *M* = 3, a central state 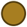 emerges, from which the action is taken almost at random. Similarly to the previous example, this state buffers the transition between the extremal ones, delaying the commitment to a particular action. For *M* = 4, this effect is ascribed to two separate central states 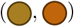 where the policy still allows taking both actions, but is skewed towards the left and right one, respectively. By marginalizing over the actions, the transitions between the internal states at fixed observation emphasize that all FSCs have a linear structure (Figure 5f). That is, the inferred internal computation *g* drives the agent towards the extremal states when a reward is obtained, whereas it transitions to the central ones in the absence of a reward. This linear structure is maintained even for larger *M*, suggesting that it can consistently model the internal decision process of the mice. The diminishing return in terms of increased accuracy from *M* = 4 onward is further highlighted by the fact that at large *M* some inferred internal states become disconnected from the rest, signaling that they are not needed to reproduce the observed behavior (see SI Appendix).

#### Emergence of internal switch detectors accurately reproduces the decision-making process

We now focus on the mechanisms that enable FSCs with larger *M* to better represent mice behavior. To this end, we generate behavioral trajectories from the FSCs for a fixed sequence of high ports (right, left, right) and track the probability of occupying a given internal state (Figure 5g-i. For *M* = 2, the FSC is able to correctly follow the sequence of environments, but the probability of occupying the state corresponding to the worst action remains high (e.g., 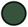 when the high port is the right one). This results in a lower probability of taking the best action, as also shown in Figure 5c.

For *M* = 3, accuracy substantially increases thanks to the middle state 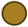. In Figure 5i we show that the probability 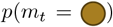 peaks right after the high port changes. Hence, 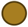 acts as a switch detector, quickly activating when the number of rewards decreases more than expected in the previous state. The FSC with *M* = 4 is able to further improve its model of behavior thanks to the emergence of two dedicated switch detectors. As we show in Figure 5i, the 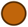 activates when the high port changes from the right to the left, whereas 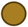 is responsible for detecting the left-to-right switch. The policies 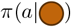 and 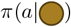 are still dominated by the right and left action, respectively, but are stochastic, allowing these states to probe the opposite action. The presence of these two switch detectors allows the accurate reconstruction of the mice behavior.

## DISCUSSION

The inference of decision-making processes hinges upon the availability of data about the interaction between agent and environment in the form of histories of actions and observations. In the cases discussed above, the identification of actions and observations is straightforward, but more complex situations command more attention. Several technical tools are now available to pre-process raw behavioral data and isolate suitable candidates for actions and observations [47–50]. In general, the key step resides in the choice of the level of description that the agent model wants to address. For instance, in a human navigation task, there is a coexistence of low-level actions and observations – like taking a step in some direction and noticing the presence of obstacles – and high-level actions and observations – like moving from one room to another and reading direction signs.

The minimalist design of agent models like FSC can prove very useful if a high-level description of behavior is sought. Indeed, an agent model based on low-level actions and observations alone typically requires a larger number of internal states to account for all the corresponding low-level computations, whereas fewer internal states often suffice at the higher level of description. Sometimes, however, it is impossible to avoid the granularity of the lowest levels of description, and this may lead to an unsustainable growth of complexity of the agent model. In these cases, a good compromise is achieved by a hierarchical organization in which different levels are stacked one upon another [51]. For instance, behavior trees [52] are a specific example of hierarchical decision structures widely used in robotics and in game development that could provide inspiration for building parsimonious agent models.

We stress once more that here we did not make any attempt at modeling the environment, nor required any prior information about it. However, in the event that some knowledge about the nature of the task is available beforehand, it can be used to appropriately design the agent model. For example, in the evidence accumulation task above, the presence of an external signal that ends the acquisition of information naturally suggests adopting a two-layer topology for the FSC. In the case of a free-response experiment in which the agent freely decides when to go right or left (see e.g. [53]), a simple linear topology turns out to be sufficient.

Importantly, FSCs can be used as generative models when a model of the environment is known. In some cases – such as the evidence accumulation tasks for rats – observations are independent of actions, and thus the inferred FSCs can generate actions in response to arbitrarily chosen sequences of observations. In general, however, the generation of behavioral trajectories requires knowledge about how observations depend on actions. In any event, using the inferred FSC in generative mode gives the possibility of predicting the agent’s response to a crafted environment and suggesting ways to steer the agent’s behavior in a desired direction.

We note that our approach considers discrete sets of actions and observations, but in real tasks these are often continuous, possibly high-dimensional variables. In these cases, it is necessary to resort to function approximation in order to represent the agent with a manageable number of parameters, e.g., by replacing the parametrization in Methods with a more general softmax with appropriately selected feature vectors. While a discussion of this approach goes beyond the scope of this paper, we can report some success in some selected tasks. Further work still remains to be done in this direction.

We focused on the setting where observations *y*_*t*_ are received by the agent after actions *a*_*t*_, representing agents that actively interact with the environment. In this case, actions can serve a dual purpose: achieving a goal – such as moving towards a target or obtaining food – and gathering information – for instance, revealing occluded objects or exploring the environment. Several, if not most, sensory processes can be described at a scale where they belong to this category [54, 55]. Other choices are possible (see Supplementary Information), and our framework can be readily applied to them as well.

The connection between FSC and neural computations is given by the formal correspondence with linear fully gated recurrent units. For the evidence accumulation tasks, this is even more explicit by the finding that the inferred FSC with *M* = 5 nodes can be interpreted as a leaky competing accumulator model. More generally, the internal states of FSC could be viewed as attractors of a recurrent neural network dynamics. This interpretation is consistent with the fact that recurrent neural networks trained on some task display a fast dynamics of convergence to attractors and slow transitions between them, which can be triggered by external inputs or occur spontaneously. This slow dynamics can be effectively represented by discrete states and transitions [24, 28, 56–58], and it has been suggested that it underlies flexible decisions and computations through dynamics [27, 59]. In this context, linear arrangements of internal states, like the ones that emerge in the analysis of rats’ and mice’s behavior, might then be interpreted as proxies for line attractors [32, 60].

In this work, we discussed applications to experimental data of rodent behavior, but our approach does not make assumptions about the nature of the agent. Any underlying computational process, be it neural, biochemical, or artificial, as long as it supports a coarse-grained description in terms of discrete states, could be a suitable candidate to be described by an FSC. It would be interesting to explore the applicability of our method to a broader range of biological agents and tasks, including human decision-making.

## I. MATERIAL AND METHODS

### FSC parametrization

A finite state controller with *M* internal states is defined by three probability distributions – the policy, the internal computation, and the initial state distribution – which we parametrize using softmax functions. The policy *π*(*a*|*m*) models the probability of taking an action *a* ∈ 𝒜 from a specific internal state *m* ∈ ℳ, with |𝒜| = *A* and |ℳ| = *M*. Hence,

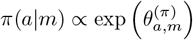

where *θ*^(*π*)^ ∈ ℝ^*A×M*^ are the parameters we seek to infer. The internal computation *g*(*m*^*′*^|*m, a, y*)models the probability of transitioning to a state *m*^*′*^ from a state *m*, conditioned on the action *a* taken by the agent and the observation *y* ∈ 𝒴 it received, with |𝒴| = *Y*. As before,

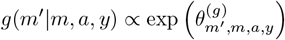

where *θ*^(*g*)^ ∈ ℝ^*M×M×A×Y*^. Finally, the initial internal state of the FSC is specified by

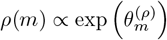

with *θ*^(*ρ*)^ ∈ ℝ^*M*^. Therefore, the total number of parameters is *M* (1 + *A* + *MAY*).

### FSC Inference

The goal is to identify the FSC which is as “close” as possible to the true unknown agent, i.e. solve the minimization problem

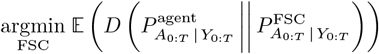

where FSC is a shorthand for the set of parameters *θ*. Above, 𝔼 is the expectation over the distribution of trajectories of actions and observations (*A*_0:*T*_, *Y*_0:*T*_) generated by the interaction of the agent and the environment, *D* is the Kullback-Leibler divergence, and

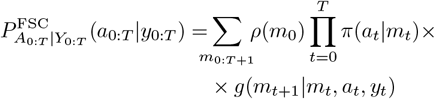

where the policy, the internal computation, and the initial distribution are parametrized as described in the previous section. Noting that

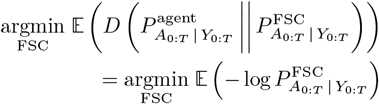

and replacing the expectation value with the empirical distribution over a set of *N* behavioral trajectories 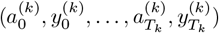 for *k* = 1, …, *N*, one arrives at the maximum-likelihood problem (1)

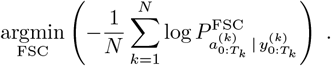

### Metric-Adaptive Particle Swarm Optimization

The main idea of MAPSO is sketched in Figure 6. First, *N*_*p*_ FSCs are initialized with random parameters. We can think of each FSC as a particle *θ*_*i*_ in the *θ* space. The parameters of each particle are updated according to a swarming dynamics, where the velocity of each particle is controlled by three different terms. First, an inertia term decreases the velocity with the number of epochs. The second term is a so-called cognitive coefficient, where the velocity vector of the *p*-th particle evolves to point towards the best set of parameters ever seen by the particle 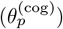. Finally, the last term is a social coefficient, where the velocity tends to point towards the best set of parameters ever seen by one of the neighbors of the particles 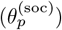, where the neighbors are specified by a network of interaction that changes as the algorithm progresses. At the beginning, during a “global phase” all particles interact with one another. Then, a “local metric phase” takes place, where at each step of the algorithm, each particle interacts only with a finite number of neighbors defined by the Euclidean distance between them. The number of neighbors decreases until the particles interact only with their nearest neighbors, before increasing back. The algorithm ends with a global phase to guarantee that all particles share the same set of best parameters.

**FIG. 6.**
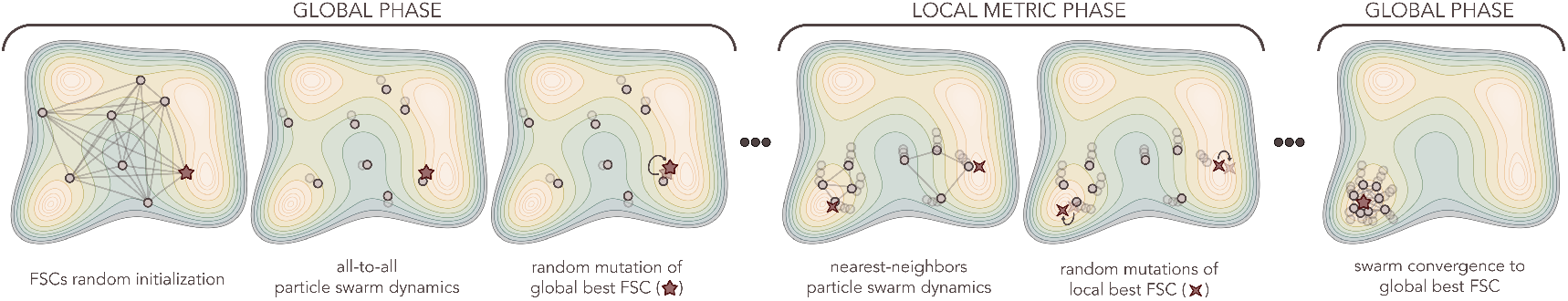
Sketch of the basic structure of metric-adaptive particle swarm optimization (MAPSO). A given number of FSCs are initialized randomly, representing a swarm of particles in the high-dimensional parameter space of the policy, computation, and initial distribution. During an initial global phase, their parameters are updated according to an all-to-all swarming dynamics, where each FSC tends to move towards the best set of parameters it has seen so far, as well as towards the best set of parameters ever found by the swarm. After each step of the swarm evolution, this global best is mutated along a random direction and is kept if the mutation improves the likelihood, and the swarm dynamics is updated according to pre-defined adaptive rules [42]. The global phase is followed by a local metric phase, where the swarm dynamics only involve a finite number of neighbors defined by the Euclidean distance between the particles. During this phase, the FSCs evolve towards the local best set of parameters, i.e., the best set of parameters ever found by the neighbors. The number of neighbors decreases before increasing back, and the algorithm ends with a second global phase.

All the parameters governing the relative strength of these terms are adaptively set according to the fuzzy rules introduced in [42]. Furthermore, at each step, the swarming dynamics is hybridized with a genetic algorithm, where each 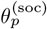 is mutated along a random direction and retained if it reduces the corresponding negative log-likelihood. MAPSO runs for a fixed number of epochs, where each epoch corresponds to a velocity update, a particle update, a parameter strength update, a computation of the metric distances between the particles and the resulting topology, and a mutation step. See the Supplementary Information for more details.

### MAPSO training schedule

To select the best inferred FSC, we consider a crossvalidation scheme. We split the data into a training set (50% of the available trajectories in the rats’ experiments, and 80% of the available trajectories in the mice’s experiments) and a validation set (the remaining trajectories). After running MAPSO for a fixed number of epochs over the training set, the best set of parameters ever found by a particle is returned. These parameters correspond to 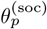 due to the final global phase where all particles interact with one another. Since the optimization problem is non-convex, we run the training procedure multiple times for a given number of internal states *M*, each with a random set of initial conditions for MAPSO. Then, we compute the negative log-likelihood over the validation set of each of the interred FSCs, and retain the one with the smallest one.

In this work, the initial parameters were extracted from a multivariate Gaussian distribution centered around zero and with diagonal covariance, but other choices are possible to bias the initial search in specific regions of the parameter space. We also note that MAPSO allows for other schedules, such as sequential restarts, where the particles’ initial position is not independent but is centered around the best parameters previously found with varying variance. See the Supplementary Information for more details.

## ACKNOWLEDGMENTS

We acknowledge useful discussions with E. Panizon and KVB Verano in the early stages of this work.

