## Supplementary information for "Decoding behavior with minimal and interpretable agent models"

### Supplementary Information: "Decoding behavior with minimal and interpretable agent model"

Giorgio Nicoletti[1] and Antonio Celani[1,2]
[1]Quantitative Life Sciences section, The Abdus Salam International Center for Theoretical Physics (ICTP), Trieste, Italy
[2]Department of Oncology, Università degli Studi di Torino, Italy

#### INFERENCE PROBLEM

We consider an agent that makes decisions in a partially observable environment. The agent takes actions $a$ according to a stochastic policy, and receives observations $y$ from the environment. Each action $a$ belongs to a finite set of elementary actions $\mathcal{A}$, and each observation $y$ to a finite set $\mathcal{Y}$. In principle, the agent's decisions may depend on the full history of previous actions, i.e., the policy is described by $P_t^{\text{agent}}(a_t \,|\, a_0, y_0, \ldots, a_{t-1}, y_{t-1})$, where the subscript $t$ is a shorthand notation to denote the conditional dependence on the history up to time $t$. Given a sequence of $T$ observations – which we do not attempt to model, since they are produced by the external environment –, the probability that the agent has taken a sequence of actions is

$$P^{\text{agent}}_{A_{0:T}|Y_{0:T}}(a_0, \ldots, a_T \,|\, y_0, \ldots, y_T) = P_0^{\text{agent}}(a_0) P_1^{\text{agent}}(a_1 \,|\, y_0, a_0) \prod_{t=1}^{T-1} P_{t+1}^{\text{agent}}(a_{t+1} \,|\, h_t) \tag{1}$$

where $h_t = (y_0, a_0, \ldots, y_t, a_t)$ is the history up to time $t$.

Then, our inference problem is the following. We seek to infer the finite state controller (FSC) that best describes the agent, where the FSC is defined by:

- a finite set of $M$ internal states $\mathcal{M} = \{M_1, \ldots, M_M\}$;
- a policy $\pi(a|m)$, describing the probability of taking an action $a \in \mathcal{A}$ from an internal state $m \in \mathcal{M}$;
- an internal computation $g(m'|m, a, y)$, describing the probability of transitioning to a new internal state $m' \in \mathcal{M}$ from a previous state $m \in \mathcal{M}$ after taking action $a \in \mathcal{A}$ and observing $y \in \mathcal{Y}$;
- an initial distribution over the internal states $\rho(m)$.

In particular, the policy is parametrized as the softmax distribution

$$\pi(a|m) = \frac{\exp\left(\theta^{(\pi)}_{a,m}\right)}{\sum_{a'} \exp\left(\theta^{(\pi)}_{a',m}\right)} \tag{2}$$

where $\theta^{(\pi)} \in \mathbb{R}^{A\times M}$ are the parameters we seek to infer, where $A = |\mathcal{A}|$ is the number of elementary actions. Similarly, the internal computation $g(m'|m, a, y)$ is

$$g(m'|m, a, y) = \frac{\exp\left(\theta^{(g)}_{m',m,a,y}\right)}{\sum_{m''} \exp\left(\theta^{(g)}_{m'',m,a,y}\right)} \tag{3}$$

where $\theta^{(g)} \in \mathbb{R}^{M\times M\times A\times Y}$, with $Y = |\mathcal{Y}|$ the number of possible observations. Finally, the initial internal state of the FSC is specified by

$$\rho(m) = \frac{\exp\left(\theta^{(\rho)}_{m}\right)}{\sum_{m'} \exp\left(\theta^{(\rho)}_{m'}\right)} \tag{4}$$

with $\theta^{(\rho)} \in \mathbb{R}^M$.

As reported in the main text, the probability that the FSC generates a sequence of actions given a sequence of observations of length $T$ can be written as

$$P^{\text{FSC}}_{A_{0:T}|Y_{0:T}} = \sum_{m_{0:T+1}} \rho(m_0) \prod_{t=0}^{T} \pi(a_t|m_t) g(m_{t+1}|m_t, a_t, y_t) \tag{5}$$

which does not depend on the internal states, i.e., $m_{0:T+1}$ can be marginalized out exactly. By definition, the FSC that best models the data is the one that minimizes the expected Kullback-Leibler divergence

$$\mathbb{E}\left[D\left(P^{\text{agent}}_{A_{0:T}|Y_{0:T}}\,||P^{\text{FSC}}_{A_{0:T}|Y_{0:T}}\right)\right]=\mathbb{E}\left[P^{\text{agent}}_{A_{0:T}|Y_{0:T}}\log\frac{P^{\text{agent}}_{A_{0:T}|Y_{0:T}}}{P^{\text{FSC}}_{A_{0:T}|Y_{0:T}}}\right] \tag{6}$$

where the expectation value is taken over the sequences of observations. We remark that both distributions must be understood in terms of their transition probabilities, i.e., of the probability of generating an action at each time $t$.

In order to infer the FSC, we collect $N$ behavioral trajectories of different durations generated by the agent, which we denote as $\{(\tilde{a}_k,\tilde{y}_k)\}_{k=1}^{N}$. Each behavioral trajectory is a sequence of actions and observations, which, for ease of notation, we denote with

$$(\tilde{a}_k,\tilde{y}_k)=\left(a_0^{(k)},y_0^{(k)},\ldots,a_{T_k}^{(k)},y_{T_k}^{(k)}\right) \tag{7}$$

where $T_k$ is the duration of the $k$-th trajectory. For ease of notation, we will also denote with $h_t=(a_0,y_0,\ldots,a_{t-1},y_{t-1})$ the history up to time $t$. From these trajectories, we can build the empirical distribution of the data,

$$\hat{P}^{\text{data}}(\tilde{a},\tilde{y})=\frac{1}{N}\sum_{k=1}^{N}\mathbb{I}\left[\tilde{y}=\left(y_0^{(k)},\ldots,y_{T_k}^{(k)}\right),\tilde{a}=\left(a_0^{(k)},\ldots,a_{T_k}^{(k)}\right)\right] \tag{8}$$

where $\mathbb{I}$ is the indicator function. Thus, replacing the expectation value in (6) with the empirical distribution, we obtain

$$\begin{aligned}\mathbb{E}\left[D\left(P^{\text{agent}}_{A_{0:T}|Y_{0:T}}\,||P^{\text{FSC}}_{A_{0:T}|Y_{0:T}}\right)\right]&\approx\sum_{\tilde{a},\tilde{y}}\hat{P}^{\text{data}}(\tilde{a},\tilde{y})\left[\log\hat{P}^{\text{data}}_{A_{0:T}|Y_{0:T}}(\tilde{a},\tilde{y})-\log P^{\text{FSC}}_{A_{0:T}|Y_{0:T}}(\tilde{a},\tilde{y})\right]\\&=\frac{1}{N}\sum_{k=1}^{N}\left[\log\hat{P}^{\text{data}}_{A_{0:T}|Y_{0:T}}(\tilde{a}_k|\tilde{y}_k)-\log P^{\text{FSC}}_{A_{0:T}|Y_{0:T}}(\tilde{a}_k|\tilde{y}_k)\right]\\&=\frac{1}{N}\sum_{k=1}^{N}\log\hat{P}^{\text{data}}_{A_{0:T}|Y_{0:T}}(\tilde{a}_k|\tilde{y}_k)+\mathcal{L}\left(\theta^{(\rho)},\theta^{(\pi)},\theta^{(g)}\right)\end{aligned} \tag{9}$$

where $\mathcal{L}$ is the negative log-likelihood of the FSC. Unrolling over the trajectories, the first term can instead be written as

$$\begin{aligned}\frac{1}{N}\sum_{k=1}^{N}\log\hat{P}^{\text{data}}_{A_{0:T}|Y_{0:T}}(\tilde{a}_k|\tilde{y}_k)&=\frac{1}{N}\sum_{k=1}^{N}\left[\log\hat{P}^{\text{data}}\left(a_0^{(k)}\right)+\sum_{t=1}^{T_k}\hat{P}^{\text{data}}\left(a_t^{(k)}\,|h_t^{(k)}\right)\right]\\&\approx H\left(A_{0:T}\,|\,Y_{0:T}\right)\end{aligned} \tag{10}$$

where $H\left(A_{0:T}\,|\,Y_{0:T}\right)$ is the entropy of the sequence of actions given the observations. We remark here that we often work with a limited number of behavioral trajectories, whereas estimating $H\left(A_{0:T}\,|\,Y_{0:T}\right)$ would require an accurate estimation of the high-dimensional full joint distribution $P^{\text{agent}}(\tilde{a},\tilde{y})$, which is in practice out of reach. Nevertheless, in some scenarios, such as the parity checker agent (see Figure 2 of the main text), the agent acts deterministically except for the first action, which is stochastic. In this case, (10) simply amounts to the entropy of the distribution of the first action, which can be reliably estimated from the data. For the parity checker agent, which takes either action "A" or "B" at random with equal probability, we thus have $H\left(A_{0:T}\,|\,Y_{0:T}\right)=\log 2$.

Finally, the inference problem requires solving

$$\underset{\text{FSC}}{\operatorname{argmin}}\ \mathbb{E}\left[D\left(P^{\text{agent}}_{A_{0:T}|Y_{0:T}}\,||P^{\text{FSC}}_{A_{0:T}|Y_{0:T}}\right)\right] \tag{11}$$

which amounts to finding the FSC parameters $\theta=\left(\theta^{(\rho)},\theta^{(\pi)},\theta^{(g)}\right)$ such that

$$\theta^*=\underset{\theta}{\operatorname{argmin}}\ \mathcal{L}\left(\theta^{(\rho)},\theta^{(\pi)},\theta^{(g)}\right) \tag{12}$$

as reported in the main text.

TABLE S1. Comparison between Finite State Controllers and Hidden Markov Models for active perception

| Model | Internal computation | Policy / Emission | |
|---|---|---|---|
| HMM | $g(m_{t+1}\|m_t)$ | $\pi(a_t\|m_t)$ | Autonomous dynamics |
| GLM-HMM | $g(m_{t+1}\|m_t)$ | $\pi(a_t\|m_t, y_{t-1})$ | Observations modulate emissions |
| IOHMM | $g(m_{t+1}\|m_t, y_t)$ | $\pi(a_t\|m_t, y_{t-1})$ | Observations affect both transitions and emissions |
| FSC | $g(m_{t+1}\|m_t, a_t, y_t)$ | $\pi(a_t\|m_t)$ | Observations and actions feed back into state transitions |

#### FINITE STATE CONTROLLERS AND STATE SPACE MODELS

##### Comparison with Hidden Markov Models

Finite state controllers may be viewed as a generalized type of state space models (SSMs) [1] with recurrent transitions. Here, we first consider the case where observations are produced by the environment following an action, as in all experimental paradigms presented in the main text. We describe this scenario as "active perception", where the agent actively takes action in order to gather information from the environment. In the literature, these have sometimes been named "information-seeking" actions and belong to the realm of active sensing [2, 3]. This is also a common setting for partially observable Markov decision processes and reinforcement learning in general, where observations can be identified with rewards and are given to the agent after an action [4].

One of the crucial features of FSCs is that transitions between internal states explicitly depend on both actions and observations. Although this recurrence makes the inference problem particularly hard, it allows them to be expressive while retaining interpretability, as we will show in the next sections. First, we briefly compare with other common SSMs (see Table S1 and Figure S1a-d). In a hidden Markov model (HMM), transitions between internal states are autonomous and depend only on the previous state ($g(m_t|m_{t-1})$), while the policy – also known as the emission probability – depends on the current internal state, as in FSCs ($\pi(a_t|m_t)$). Thus, observations do not influence transitions. A possible generalization is HMMs where emissions are parametrized by a generalized linear model (GLM-HMMs), so that the policy explicitly depends on observations [5]. More generally, in input-output hidden Markov models (IOHMMs), the dependence on observations enters both the emission probability and the internal computation [6]. However, the basic idea of these state space models is that the dynamics of the internal or hidden states is disentangled from that of the emissions, which is instead a key dependency in FSCs.

##### Moore and Mealy formulations of FSCs and dependence on past observations

In frameworks for describing behavior and decision-making, policies can be formulated in two ways: they can depend only on the internal state – a Moore formulation –, or they can depend on both the internal state and external observations – a Mealy formulation [4, 7]. As formulated in the main text and so far, FSCs implement a Moore architecture where the policy $\pi(a_t|m_t)$ depends only on the current internal state. In contrast, IOHMMs follow a Mealy machine structure with observation-dependent emissions, which, for the case of active perception, gives $\pi(a_t|m_t, y_{t-1})$ to preserve the causal structure. Nevertheless, FSCs achieve expressiveness through recurrent state updates $g(m_{t+1}|m_t, a_t, y_t)$ rather than observation-dependent policies. That is, since the history $h_t = (a_0, y_0, \ldots, a_{t-1}, y_{t-1})$ is encoded in the internal states, the policy of an FSC implicitly depends on both past actions and observations, allowing them to exhibit history-dependent behavior.

More rigorously, we can always choose a structure of internal states that makes the dependence on the previous observation explicit. One could formulate an equivalent Mealy-type FSC by augmenting the internal state space to explicitly include the previous observation, e.g., $m_t = (\tilde{m}_t, \hat{m}_t)$ where

$$g(m_{t+1}|m_t, a_t, y_t) = g(\tilde{m}_{t+1}|m_t, a_t, y_t)\delta(\hat{m}_{t+1}, y_t) \tag{13}$$

so that $\hat{m}_t$ perfectly remembers the previous observation. The policy would then be $\pi(a_t|\tilde{m}_t, y_{t-1})$, making the dependence on recent observations explicit rather than implicit. While both formulations are mathematically equivalent and equally expressive, the Moore structure offers a more compact state representation, requiring $|\mathcal{M}|$ states instead of $|\mathcal{M}| \times |\mathcal{Y}|$. Furthermore, the Moore formulation provides greater interpretability: each internal state has a well-defined behavioral signature – the actions it produces–, while state transitions capture how the FSC updates its internal representation in response to past observations and actions. This separation between action selection and state update proves particularly useful for understanding the inferred FSCs, as we show in the main text and in later sections.

##### Active perception and stimulus-driven FSCs

Even though we have focused on the case of active perception, both FSCs and our inference method can be formulated to account for scenarios that are instead "stimulus-driven", i.e., where observations are received by the agent before taking actions [8, 9]. As we show in Figure S1e, in this case, the natural formulation of an FSC is in the form of a Mealy machine, i.e.,

$$\pi(a_t|m_t, y_t), \quad g(m_{t+1}|m_t, a_t, y_t) \ . \tag{14}$$

We stress that the distinction between active perception and stimulus-driven behaviors strictly depends on the causal structure of the environment, which we do not observe nor model since the likelihood in (5) is completely independent of the environment itself. We also highlight that the dynamics of the environment could be, in principle, extremely complex – e.g., actions taken at a given time may have long-term influences on the observations received by the agent. Furthermore, the behavioral trajectories themselves or the experimental design should determine whether an active perception or a stimulus-driven description is more appropriate. Arguably, one can always describe the decision-making process at a scale where it falls under the active perception framework [2, 10]. For instance, the agent may choose to take an action "perceive" or "sense" to passively receive observations from the environment. Clearly, this may drastically increase the space of possible actions and observations, making a stimulus-driven description more useful in practice. In Figure S1f-h, we compare this FSC with the corresponding HMMs, showing once more how it generalizes their structure by including recurrent transitions and allowing the agent to encode past decisions in its internal states.

#### INTERPRETING FSCS' INTERNAL COMPUTATIONS

##### Behaviorally-equivalent FSC and invariance under linear mixing

Before showing how the internal structure of finite state controllers can be explicitly interpreted in terms of behavioral modes and internal computations, we show here that in some cases two different FSCs may sometimes lead to the same behavior. More precisely, two FSCs described by initial distributions $\rho$ and $\rho'$, policies $\pi$ and $\pi'$, and internal computations $g$ and $g'$ are behaviorally equivalent if they generate the same conditional probabilities for all histories of actions and observations, i.e., $p(a_{t+1}|h_t) = p'(a_{t+1}|h_t)$ with $h_t = (y_0, a_0, \dots, y_t, a_t)$ as defined above.

In particular, for an FSC with $M$ internal states, we define the observable transition (OT) matrices $(O_{a,y}) \in \mathbb{R}_{\geq 0}^{M\times M}$ as the matrices whose elements $(O_{a,y})_{m'm}$ are equal to the joint transition probabilities

$$O(m', a|m, y) = \pi(a|m) g(m'|m, a, y) \tag{15}$$

which fully describes the FSC, since (15) is nothing but the definition of conditional probability. Then, we seek an invertible matrix $T \in \mathbb{R}_{\geq 0}^{M\times M}$ that leaves the action of the policy on any probability vector invariant, i.e.,

$$\Pi' T = \Pi \tag{16}$$
$$1^T T = 1^T \tag{17}$$

where we denoted with $\Pi \in \mathbb{R}_{\geq 0}^{A\times M}$ the matrix whose entries are the FSC policy. The last condition in (16) implies that $T$ is column-stochastic, so that $T$ represents a convex linear mixing of the internal states. Then, the two FSCs are equivalent if

$$\rho' = T\rho \tag{18}$$
$$O'_{a,y} = T O_{a,y} T^{-1} \quad \forall a, y \tag{19}$$
$$\tag{20}$$

where $p_{h_t} \in \Delta^A$ is the $A$-dimensional vector whose entries are $p(a_{t+1}|h_t)$, and $\Delta$ denotes the simplex. Indeed, if such a $T$ exists, we have that

$$\begin{aligned} p'_{h_t} &= \pi'_{a_{t+1}} O'_{a_t,y_t} \dots O'_{a_1,y_1} O'_{a_0,y_0} \rho' \\ &= \pi_{a_{t+1}} T^{-1} T O_{a_t,y_t} T^{-1} \dots T O_{a_1,y_1} T^{-1} T O_{a_0,y_0} T^{-1} T \rho \\ &= p_{h_t} \end{aligned} \tag{21}$$

so that the two FSCs produce statistically identical behavioral trajectories.

We remark here that, most often, $T$ is trivial. Indeed, if at least two $O_{a,y}$ are full-rank, then $T$ can only be a permutation matrix. Indeed, if $T$ were not a permutation matrix, then some column $t_j$ would have at least two non-zero entries – meaning that state $j$ in one FSC maps to a convex combination of the states of the other FSC, which is incompatible with the similarity on the OT matrices, $TO'_{a,y} = TO_{a,y}$ (see [11] for more details). However, when at least $AY - 1$ OT matrices are not full-rank – i.e., there is at most one OT matrix whose rank is $M$ – then $T$ can be different from a permutation. In the chemotactic agent presented in the main text, this is the case. In Figure S2, we show two FSCs that can both be obtained from the inference procedure, both with the same negative log-likelihood – and equal to that of the original chemotactic agent – that are described by

$$\Pi_1 = \begin{pmatrix} 1/2 & 1 & 0 & 1/2 \\ 1/2 & 0 & 1 & 1/2 \end{pmatrix}, \quad \Pi_2 = \begin{pmatrix} 1/2 & 1 & 0 & 0 \\ 1/2 & 0 & 1 & 1 \end{pmatrix} \tag{22}$$

where the first row is run ($R$), the second tumble ($T$). The internal computations for the first FSC, the one presented in the main text, are given by

$$G_1(y=\bullet, a=R) = \begin{pmatrix} 0 & 0 & ? & 0 \\ 1 & 0 & ? & 0 \\ 0 & 0 & ? & 0 \\ 0 & 1 & ? & 1 \end{pmatrix}, \quad G_1(y=\times, a=R) = \begin{pmatrix} 1 & 0 & ? & 0 \\ 0 & 0 & ? & 0 \\ 0 & 1 & ? & 1 \\ 0 & 0 & ? & 0 \end{pmatrix} \tag{23}$$

$$G_1(y=\bullet, a=T) = \begin{pmatrix} 0 & ? & 0 & 0 \\ 1 & ? & 1 & 0 \\ 0 & ? & 0 & 0 \\ 0 & ? & 0 & 1 \end{pmatrix}, \quad G_1(y=\times, a=T) = \begin{pmatrix} 1 & ? & 1 & 0 \\ 0 & ? & 0 & 0 \\ 0 & ? & 0 & 1 \\ 0 & ? & 0 & 0 \end{pmatrix} \tag{24}$$

where $G(y,a)$ are the matrices whose elements $G(y,a)_{m'm}$ are equal to the internal computation $g(m'|a,y,m)$, the question marks denote undefined transitions (because the policy does not allow the action from the specific state), and $\{\bullet, \times\}$ represent observing or not observing a chemical, respectively. The second FSC, instead, is described by

$$G_2(y=\bullet, a=R) = \begin{pmatrix} 0 & 0 & ? & ? \\ 1 & 1/2 & ? & ? \\ 0 & 0 & ? & ? \\ 0 & 1/2 & ? & ? \end{pmatrix}, \quad G_2(y=\times, a=R) = \begin{pmatrix} 1 & 0 & ? & ? \\ 0 & 0 & ? & ? \\ 0 & 1 & ? & ? \\ 0 & 0 & ? & ? \end{pmatrix} \tag{25}$$

$$G_2(y=\bullet, a=T) = \begin{pmatrix} 0 & ? & 0 & 0 \\ 1 & ? & 1 & 1/2 \\ 0 & ? & 0 & 0 \\ 0 & ? & 0 & 1/2 \end{pmatrix}, \quad G_2(y=\times, a=T) = \begin{pmatrix} 1 & ? & 1 & 0 \\ 0 & ? & 0 & 0 \\ 0 & ? & 0 & 1 \\ 0 & ? & 0 & 0 \end{pmatrix} \tag{26}$$

and both are initialized in their respective first state.

Consider the matrix

$$T = \begin{pmatrix} 1 & 0 & 0 & 0 \\ 0 & 1 & 0 & 1/2 \\ 0 & 0 & 1 & 0 \\ 0 & 0 & 0 & 1/2 \end{pmatrix} \tag{27}$$

which is column-stochastic and ensures that

$$\pi_2 T = \pi_1 \tag{28}$$

and satisfies trivially $\rho_2 = T\rho_1$. It is then easy to show that the observable transition matrices,

$$O_1(y=\bullet, a=R) = \begin{pmatrix} 0 & 0 & 0 & 0 \\ 1/2 & 0 & 0 & 0 \\ 0 & 0 & 0 & 0 \\ 0 & 1 & 0 & 1/2 \end{pmatrix}, \quad O_1(y=\times, a=R) = \begin{pmatrix} 1/2 & 0 & 0 & 0 \\ 0 & 0 & 0 & 0 \\ 0 & 1 & 0 & 1/2 \\ 0 & 0 & 0 & 0 \end{pmatrix} \tag{29}$$

$$O_1(y=\bullet, a=T) = \begin{pmatrix} 0 & 0 & 0 & 0 \\ 1/2 & 0 & 1 & 0 \\ 0 & 0 & 0 & 0 \\ 0 & 0 & 0 & 1/2 \end{pmatrix}, \quad O_1(y=\times, a=T) = \begin{pmatrix} 1/2 & 0 & 1 & 0 \\ 0 & 0 & 0 & 0 \\ 0 & 0 & 0 & 1/2 \\ 0 & 0 & 0 & 0 \end{pmatrix} \tag{30}$$

and

$$O_2(y=\bullet, a=R) = \begin{pmatrix} 0 & 0 & 0 & 0 \\ 1/2 & 1/2 & 0 & 0 \\ 0 & 0 & 0 & 0 \\ 0 & 1/2 & 0 & 0 \end{pmatrix}, \quad O_2(y=\times, a=R) = \begin{pmatrix} 1/2 & 0 & 0 & 0 \\ 0 & 0 & 0 & 0 \\ 0 & 1 & 0 & 0 \\ 0 & 0 & 0 & 0 \end{pmatrix} \tag{31}$$

$$O_2(y=\bullet, a=T) = \begin{pmatrix} 0 & 0 & 0 & 0 \\ 1/2 & 0 & 1 & 1/2 \\ 0 & 0 & 0 & 0 \\ 0 & 0 & 0 & 1/2 \end{pmatrix}, \quad O_2(y=\times, a=T) = \begin{pmatrix} 1/2 & 0 & 1 & 0 \\ 0 & 0 & 0 & 0 \\ 0 & 0 & 0 & 1 \\ 0 & 0 & 0 & 0 \end{pmatrix} \tag{32}$$

are all of rank $2 < M = 4$, and satisfy $O_2 = TO_1T^{-1}$ for all $a, y$, making the two FSC behaviorally equivalent, as expected – since they both reach the negative log-likelihood of the original agent. We also note here that any FSC is properly described in terms of the observable transition matrices alone. For example, in these two cases, some internal computations are not defined because the policy does not allow taking the corresponding action from the corresponding internal state. Because of this, OT matrices are always well-defined.

#### FSCs with two internal states

As shown in the main text, finite state controllers are expressive enough to reproduce a wide range of behaviors. Importantly, at least in some cases, their finite number of internal states allows for a direct interpretation of their internal computations. Here, we focus on the simplest case of $M = 2$, where we can make general statements. Consider, as above, the $M \times M$ matrix $G(a, y)$ defined by the internal computation $g(m'|m, a, y)$. We denote with $q(a, y)$ the steady state of $G(a, y)$, i.e., the eigenvector of eigenvalue 1 of $G(a, y)$, such that $G(a, y)q(a, y) = q(a, y)$. Similarly, $r(a, y)$ and $l(a, y)$ are the right and left eigenvectors of eigenvalue $\lambda_2$ of $G(a, y)$. Then, we have that

$$\begin{aligned} G(a,y) &= q(a,y)1^T + |\lambda_2(a,y)|\, r(a,y)\, l(a,y)^T \\ &= [1 - |\lambda_2(a,y)|]\, q(a,y)1^T + |\lambda_2(a,y)| \left[ r(a,y)l(a,y)^T + q(a,y)1^T \right] \\ &= |\lambda_2(a,y)|\mathbb{I} + (1 - |\lambda_2(a,y)|)q(a,y)1^T \end{aligned} \tag{33}$$

where 1 if the vector of all ones and $\mathbb{I}$ is the identity matrix. Therefore, if we now write the probability of occupying an internal state at time $t$ as $p_{\mathrm{mem}}(t) = (p_1(t), p_2(t))$, with $p_2(t) = 1 - p_1(t)$, after taking action $a_t$ and observing $y_t$ this probability evolves as

$$p_1(t+1) = |\lambda_2(a_t, y_t)|p_1(t) + [1 - |\lambda_2(a_t, y_t)|]\, q_1(a_t, y_t) \tag{34}$$

where we have written the components of the stationary state as $q = (q_1, 1 - q_1)$. Therefore, the internal state of the FSC associated with $\phi$ is computing a discounted sum of the action-observation pairs $(a, y)$, with a discount factor $\lambda_2(a, y) < 1$ that depends on the pair itself. We can interpret this first term in (34) as a forgetting term that manifestly depends on the relaxation timescale of $G(a, y)$. The internal representation of each pair is instead the corresponding stationary state of the dynamics at fixed $(a, y)$, allowing the FSC to encode different pairs in different points of the simplex. Thus, this internal representation effectively depends on how different the stationary states are.

In the case of the computation layer of the FSC with $M = 4$ for the evidence accumulation tasks, for the listen action, we have

$$G(y) = \begin{pmatrix} 1 - \epsilon y \frac{y-1}{2} & \epsilon y \frac{1+y}{2} \\ \epsilon y \frac{y-1}{2} & 1 - \epsilon y \frac{1+y}{2} \end{pmatrix}, \qquad q(y) = \frac{1}{2} \begin{pmatrix} 1+y \\ 1-y \end{pmatrix}, \qquad \lambda_2(y) = 1 - \epsilon y^2 \tag{35}$$

where $y \in \{0, \pm 1\}$ for the silence, right, and left observations, and $\epsilon$ is the probability of switching from one memory to the other if a contrasting observation is received. Therefore, as in the main text, if we define $\phi(t) = p_1(t) - p_2(t) = 2p_1(t) - 1$, we have that

$$\phi(t+1) = \lambda_2(a_t, y_t)\phi(t) + [1 - \lambda_2(a_t, y_t)]\,[2q_1(a_t, y_t) - 1] \tag{36}$$

which becomes

$$\phi(t+1) = (1 - \epsilon y_t^2)\phi(t) + \epsilon y_t \tag{37}$$

so that the dynamics is fully determined by $\epsilon$. If $\epsilon = 1$, the FSC becomes purely reactive and implements a win-stay-lose-shift dynamics. We also note that, in this simple case where $G$ depends only on $y$, the two internal states are directly encoding for the left ($y = -1$) or right ($y = 1$) observations, since the silence ($y = 0$) does not change the internal state as expected.

##### FSCs with an arbitrary number of internal states

Let us define the probability that an FSC with $M$ internal states occupies an internal state at time $t$ as the belief state $b_t \in \Delta^M$, where $\Delta^M$ is the $M$-dimensional simplex. The system receives a discrete input tuple $(a_t, y_t)$ at each time step. As before, we map each unique pair $(a, y)$ to a unique index $k \in \{1, \ldots, K\}$. Then, the belief update is

$$b_{t+1} = G_k b_t \tag{38}$$

where the $k$ index depends on time. We decompose $G_k$ into its stationary and transient components as

$$G_k = \tilde{G}_k + (1 - \gamma_k) q_k 1^T \tag{39}$$

where $\gamma_k = |\lambda_2(G_k)|$ is the modulus of the second largest eigenvalue modulus satisfying $|\lambda_2(G_k)| < 1$, and as before $q_k$ is the steady-state distribution of $G_k$, i.e., $G_k q_k = q_k$. We further decompose the matrix $\tilde{G}_k$ as

$$\tilde{G}_k = H_k + \gamma_k \mathbb{I} \tag{40}$$

so that, since $1^T b_t = 1$, we can write

$$b_{t+1} = \underbrace{\gamma_k b_t}_{\text{Decay}} + \underbrace{(1 - \gamma_k) q_k}_{\text{Injection}} + \underbrace{H_k b_t}_{\text{Mixing}} \;. \tag{41}$$

Although in general $H_k$ is at most of rank $M - 1$, if $\lambda_2(G_k)$ is real and positive, its rank is $M - 2$ since it has two zero eigenvalues – one for the steady-state, and one for the second mode. Indeed, for $M = 2$, the mixing term is not present and we recover the case of the previous section.

The first term in (41) represents the memory persistence. The smaller $\gamma_k$, the shorter the memory persistence – in the case of $M = 2$, this term can be directly interpreted as the exit probability from the internal states. The second term, instead, injects back into the dynamics the steady-state of the current action-observation pair $k$, with an amplitude that is proportional to the leakage from the current belief $b_t$, i.e., $1 - \gamma_k$ . Finally, the last term mixes the different beliefs together, adding or subtracting probability to each internal state depending on the observations and actions. In general, if $\gamma_k \approx 1$ the process is slow-mixing and the FSC tends to preserve memory. If $\gamma_k \approx 0$, the FSC favors new evidence and the process is fast-mixing. Notably, (41) is analogous to a linearized version of a Gated Recurrent Unit (GRU) [12] where the embedding of the $k$-th action-observation pair is the steady-state of $G_k$, and without a reset gate (as in light gated recurrent units [13]):

$$b_{t+1} = \gamma_k b_t + (1 - \gamma_k)\left[q_k + \frac{1}{1 - \gamma_k} H_k b_t\right] \tag{42}$$

where the term in the square bracket is the candidate belief, in analogy with the candidate state of GRUs. This form of the dynamics comes from the fact that the mixing term is originally ungated, so that when $\gamma_k \to 1$ the embedded input $q_k$ is ignored in favor of internal mixing. More detailed interpretations depend on the specific form of $G_k$ inferred from data.

**FSC with $M = 5$ inferred from evidence accumulation data**

We now focus on the FSC with $M = 5$ internal states inferred from rats' evidence accumulation data. In particular, we consider the three internal states belonging to the computation layer, since they are responsible for integrating sensory cues – the two remaining states simply project the results of the computation layer into a policy which chooses either right or left. Thus, the components of $b_t$ represent the probability of occupying 🟢, 🟡, 🔴 respectively. Using the notation of the previous sections, we consider for simplicity the perfectly symmetric FSC defined by the transition matrices

$$G(1) = \begin{pmatrix} 1-\xi & 0 & 0 \\ \xi & 1-\eta & 0 \\ 0 & \eta & 1 \end{pmatrix}, \quad G(-1) = \begin{pmatrix} 1 & \eta & 0 \\ 0 & 1-\eta & \xi \\ 0 & 0 & 1-\xi \end{pmatrix} \tag{43}$$

where $G_k = G(y_k)$, since the action is always "listen" in this case, $y = \pm 1$ correspond to hearing right or left, respectively, and $\xi, \eta \in (0, 1)$. We ignore the case of $y_t = 0$, since $G(0)$ is simply the identity matrix and no information is present in the observation. Although the inferred FSC is slightly asymmetric due to the noise in the data and the intrinsic asymmetries in the rats' behavior ($p($🟡$|$🟢$, y = 1) \approx 0.12$ and $p($🟡$|$🔴$, y = -1) \approx 0.15$, $p($🔴$|$🟡$, y = 1) \approx 0.25$ and $p($🟢$|$🟡$, y = -1) \approx 0.29$), it can be numerically checked that this remains an excellent approximation in terms on negative log-likelihood if we use the transition matrices above with average parameters $\xi \approx 0.13$ and $\eta \approx 0.27$. Similarly, the policy can be approximately expressed in terms of these three internal states as

$$\Pi = \begin{pmatrix} 1 & 1/2 & 0 \\ 0 & 1/2 & 1 \end{pmatrix} \tag{44}$$

since, when the "end" observation is received, the states of the computation layer transition to the ones in the second layer, which deterministically choose the action. The first row of $\Pi$ corresponds to the action "go left", the second to "go right".

Since $\xi < \eta$, for both $G(1)$ and $G(-1)$ we have $\lambda_2 = 1 - \xi$, whereas the stationary states are simply

$$q(1) = \begin{pmatrix} 0 \\ 0 \\ 1 \end{pmatrix}, \quad q(-1) = \begin{pmatrix} 1 \\ 0 \\ 0 \end{pmatrix} \tag{45}$$

which corresponds to the trivial fact that when only $y = 1$ is heard the FSC occupies the 🔴 state, and 🟢 if it receives only $y = -1$ instead. In this case, $H(y)$ is a rank-1 matrix, so that

$$G(y) = (1 - \xi)I + \xi q(y)1^T + u(y)v(y)^T \tag{46}$$

where

$$u(1) = \begin{pmatrix} 0 \\ 1 \\ -1 \end{pmatrix} \quad (\text{flow 🔴} \rightarrow \text{🟡}), \quad u(-1) = \begin{pmatrix} 1 \\ -1 \\ 0 \end{pmatrix} \quad (\text{flow 🟡} \rightarrow \text{🟢}) \tag{47}$$

represent the flow between the internal states, and

$$v(1) = \begin{pmatrix} \xi \\ \xi - \eta \\ 0 \end{pmatrix}, \quad v(-1) = \begin{pmatrix} 0 \\ \eta - \xi \\ -\xi \end{pmatrix} \tag{48}$$

are the corresponding amplitudes. To make the FSC dynamics more explicit, we define two scalar state variables, $\phi_t^+$ and $\phi_t^-$, as the projections:

$$\phi_t^+ := -v(-1)^T b_t \tag{49}$$

$$\phi_t^- := v(1)^T b_t \tag{50}$$

where $\phi_t^+$ is a positive accumulator representing evidence favoring state 🔴 ($y = 1$), and $\phi_t^-$ a negative accumulator representing evidence favoring state 🟢 ($y = -1$). The dynamics of the pair $(\phi_t^+, \phi_t^-)$ forms a switched linear system with asymmetric lateral inhibition. We distinguish two cases. With a positive input ($y_t = 1$) the positive accumulator is *active* (driven), while the negative accumulator is *passive* (decaying):

$$\phi_t^+ = (1-\xi)\phi_{t-1}^+ - \eta\phi_{t-1}^- + \xi^2 \tag{51}$$
$$\phi_t^- = (1-\eta)\phi_{t-1}^- \tag{52}$$

so that the passive unit inhibits the active unit. With a negative input ($y_k = -1$), the roles are switched (the negative accumulator is active, and the positive accumulator is passive):

$$\phi_t^+ = (1-\eta)\phi_{t-1}^+ \tag{53}$$
$$\phi_t^- = (1-\xi)\phi_{t-1}^- - \eta\phi_{t-1}^+ + \xi^2 \tag{54}$$

and the inhibition direction flips. Crucially, this is a discrete version of a leaky competing accumulator model [14] (LCA) with:

1. accumulation – the active variable integrates the evidence source with a term $\xi^2$;
2. forgetting – the active unit leaks at rate $\xi$, the passive unit at a rate $\eta$;
3. switched lateral inhibition – the passive unit inhibits the active unit via the term $-\eta\phi_{t-1}^{\mp}$, but not vice versa.

Thus, when the input switches, the new target accumulator is initially suppressed by the inhibition due to the stored value of the old accumulator, generating a dynamics where previous strong commitments are more persistent, as shown in the main text. We also note that the passive decay $(1-\eta)$ ensures that the inhibition eventually vanishes, allowing the active unit to reach its steady state in the presence of a fully coherent sequence of observations.

In terms of the internal states of the FSC, the accumulators are given by

$$\phi_t^+ = (\xi-\eta)p_t({\color{goldenrod}\bullet}) + \xi p_t({\color{darkred}\bullet}) \tag{55}$$
$$\phi_t^- = \xi p_t({\color{darkgreen}\bullet}) + (\xi-\eta)p_t({\color{goldenrod}\bullet}) \tag{56}$$

where $p_t(\cdot)$ are the components of $b_t$ at time $t$, and depend on all the previous observations $y_{0:t-1}$, but we dropped the conditional dependency for brevity. Thus, we can express the probability of choosing an action over the other at the end of the trial at time $T$ as

$$\phi(T+1) = p(A_{T+1} = \text{go right}|y_{0:T}) - p(A_{T+1} = \text{go left}|y_{0:T}) \tag{57}$$
$$= \begin{pmatrix}-1 & 1\end{pmatrix} \Pi\, b_T = p_T({\color{darkred}\bullet}) + \frac{1}{2}p_T({\color{goldenrod}\bullet}) - \left(\frac{1}{2}p_T({\color{goldenrod}\bullet}) + p_T({\color{darkred}\bullet})\right) \tag{58}$$
$$= p_T({\color{darkred}\bullet}) - p_T({\color{darkgreen}\bullet}) = \frac{1}{\xi}\left(\phi_T^+ - \phi_T^-\right) \tag{59}$$

where $y_T = \text{end}$. Thus, the final decision is taken as a function of the accumulators, which accumulate observations and compete with each other via inhibition.

#### METRIC-ADAPTIVE PARTICLE SWARM OPTIMIZATION (MAPSO)

MAPSO is a global optimization algorithm based on an adaptive version of particle swarm optimization [15]. Suppose that we have a loss function to minimize, $\mathcal{L}(\theta)$, that depends on some $D$-dimensional parameters $\theta \in \mathbb{R}^D$. In the case described in the main text, $\mathcal{L}$ is the negative log-likelihood, and $\theta = (\theta^{(\pi)}, \theta^{(g)}, \theta^{(\rho)})$ are the parameters specifying the policy, computation, and initial distribution of an FSC. If $A$ is the number of actions, $Y$ the number of observations, and $M$ the number of internal states of the FSC, then $D = MA(1+MY)+M$. In general, however, MAPSO can be used with any kind of loss function – e.g., we may include constraints on specific observables that we can write as functions of $\theta$ or sparsity constraints.

The core idea of MAPSO is to explore the parameter space by evolving the position of $N_p$ particles, each corresponding to a set of parameters $\theta_\mu$ for $\mu = 1,\dots,N_p$. The particle interacts with its neighbors through a temporal interaction network $A_{\mu\nu}(t)$ that depends on all particles' positions in the parameter space. More in detail, the $i$-th component of the $\mu$-th particle is updated according to the swarming dynamics

$$\begin{aligned}\theta_{\mu,i}(t+1) &= \theta_{\mu,i}(t) + v_{\mu,i}(t+1)\\ v_{\mu,i}(t+1) &= w(t)v_{\mu,i}(t) + c_{\text{cog}}(t)\left[\theta_{\mu,i}^{(\text{cog})}(t) - \theta_{\mu,i}(t)\right] + c_{\text{soc}}(t)\left[\theta_{\mu,i}^{(\text{soc})}(t) - \theta_{\mu,i}(t)\right]\end{aligned} \tag{60}$$

for $i = 1,\dots,D$ and where:

- $w(t)$ is an adaptive inertial parameter;
- $\theta_\mu^{(\text{cog})}(t)$ is the best parameters set ever saw by the $\mu$-th particle after $t$ steps, i.e., $\theta_\mu^{(\text{cog})}(t) = \text{argmin}_\theta \left\{\mathcal{L}(\theta_\mu(\tau))\right\}_{\tau=1,\ldots,t}$, also known as the "personal best" of the $\mu$-th particle;
- $c_{\text{cog}}(t)$ is an adaptive coefficient that regulates the strength of the "cognitive" dynamics, which makes the particle move in the direction of its personal best;
- $\theta_\mu^{(\text{soc})}(t)$ is the best parameters set ever saw by the *neighbors* of the $\mu$-th particle after $t$ steps, defined at each time according to the corresponding interaction graph $A_{\mu\nu}$, an it is also known as the "local best" of the $\mu$-th particle;
- $c_{\text{soc}}(t)$ is an adaptive coefficient that regulates the strength of the "social" dynamics, which makes the particle move in the direction of its local best.

Finally, depending on the particles' positions, MAPSO attempts to mutate the local best to improve its location. Although different training schedules are possible, we now focus on the case where the algorithm runs for a fixed number of epochs $N_{\text{epochs}}$, and returns the local best with the lowest loss. We will now outline the main steps of MAPSO, and provide a pseudo-code in Algorithm 1.

##### Initialization

First, the $N_p$ particles are randomly initialized. In our case, we take each initial parameter to follow an independent Gaussian distribution, i.e., $\theta_{\mu,i}(0) \sim \mathcal{N}(0, \sigma^2)$. Similarly, the velocities of the particles are initialized from a uniform distribution $v_{\mu,i}(0) \sim U(v_{\min}, v_{\max})$. We usually take $\sigma^2 \in [0.5, 1, 2.5]$, depending on the problem, and $v_{\max} = -v_{\min} = 10^{-2}$. In general, however, any initialization is possible in principle and, in our experience, will affect the optimization results.

For each particle, the personal best is initialized by definition to the particle's initial position, $\theta_\mu^{(\text{cog})}(0) = \theta_\mu(0)$. We also set $A_{\mu\nu}(0) = 1$ for all $\mu \neq \nu$ and compute the particle for which the loss is lowest, i.e., $\mu^* = \text{argmin}_\mu \left[\mathcal{L}(\theta_\mu(0))\right]$. Since all particles initially interact with one another, each particle's local best is $\theta_\mu^{(\text{soc})} = \theta_{\mu^*}$. Following Ref. [15], we set $w(0) = 0.9$ and $c_{\text{soc}}(0) = c_{\text{cog}}(0) = 2$.

##### Personal and local best update

After a swarming dynamics step, described in (60), each particle attempts to update both its personal and its local best. We have

$$\theta_\mu^{(\text{cog})}(t+1) = \underset{\theta}{\text{argmin}} \left\{\mathcal{L}\left(\theta_\mu(t+1)\right), \mathcal{L}\left(\theta_\mu^{(\text{cog})}(t)\right)\right\} \tag{61}$$

so that the personal best is updated with the current position if it has a lower loss. Similarly, the local best is

$$\theta_\mu^{(\text{soc})}(t+1) = \underset{\theta}{\text{argmin}} \left\{\left\{\mathcal{L}\left(\theta_\nu^{(\text{cog})}(t+1)\right) \mid A_{\mu\nu} \neq 0\right\}_{\nu=1,\ldots,N_p}, \mathcal{L}\left(\theta_\mu^{(\text{soc})}(t)\right)\right\} \tag{62}$$

so that it is updated if any of the neighbors of the $\mu$-th particle has found a parameter set with a loss lower than the one of the current local best.

##### Metric interaction network update

The interaction network changes in time according to the metric distance between the particles. Heuristically, MAPSO starts with a "global phase" all particles interact with one another, then evolves towards a "local metric phase", where each particle interacts only with a finite number of neighbors, before moving back to a global phase. In practice, at each step, we compute a number of neighbors $n_{\text{neigh}}(t)$ as

$$n_{\text{neigh}}(t) = \begin{cases} (N_p - 1) + (n_{\text{neigh,min}} - (N_p - 1)) \left(\frac{t}{N_{\text{epochs}}}\right)^4 & \text{if} \quad t < \frac{N_{\text{epochs}}}{2} \\ n_{\text{neigh,min}} + (N_p - 1 - n_{\text{neigh,min}}) \left(\frac{2t - N_{\text{epochs}}}{N_{\text{epochs}}}\right)^4 & \text{if} \quad t < \frac{N_{\text{epochs}}}{2} \end{cases} \tag{63}$$

which guarantees that at the beginning and at the end the number of neighbors is equal to $N_p - 1$, while also allowing for an initial slow decay. We empirically found that similar functional forms for $n_{\text{neigh}}(t)$ do not affect the results, as long as they respect these requests.

The interaction network is then a $n_{\text{neigh}}(t)$-nearest neighbors graph, based upon the Euclidean metric. At each step, sfter determining $n_{\text{neigh}}(t)$, we compute the distance matrix

$$d_{\mu\nu}(t) = \sum_{i=1}^{D} \sqrt{\left[\theta_{\mu,i}(t) - \theta_{\nu,i}(t)\right]^2} \tag{64}$$

and define the interaction network as

$$A_{\mu\nu}(t) = \mathbb{I}\left(d_{\mu\nu}(t) < d_{\mu}^{(n_{\text{neigh}})}(t)\right) \tag{65}$$

where $d_{\mu}^{(n_{\text{neigh}})}(t)$ denotes the $n_{\text{neigh}}(t)$-th smallest values among $\{d_{\mu\nu'}(t) \,|\, \nu' \neq \mu\}$, and we follow the usual convention in graph theory that $A_{\mu\nu}$ defines the interaction from $\mu$ to $\nu$. Since the network is undirected by definition, $A_{\mu\nu} = A_{\nu\mu}$.

##### Adaptive parameters update

The parameters are updated according to the fuzzy rules introduced in Ref. [15]. In particular, an evolutionary parameter $f(t)$ is introduced, defined as

$$f(t) = \frac{d_{\text{best}}(t) - d_{\text{min}}(t)}{d_{\text{max}}(t) - d_{\text{min}}(t)} \in [0, 1] \tag{66}$$

where

$$d_{\text{min}}(t) = \min_{\mu} \frac{1}{N_p - 1} \sum_{\nu=1}^{N_p} d_{\mu\nu}(t), \quad d_{\text{max}}(t) = \max_{\mu} \frac{1}{N_p - 1} \sum_{\nu=1}^{N_p} d_{\mu\nu}(t), \quad d_{\text{best}}(t) = \frac{1}{N_p - 1} \sum_{\nu=1}^{N_p} d_{\mu^*(t)\nu}(t) \tag{67}$$

with $\mu^*(t) = \text{argmin}_{\mu} \theta_{\mu}^{(\text{soc})}(t)$. From $f(t)$, we define a membership values $S$ for four different strategies:

$$S_1(f) = \begin{cases} 0 & \text{if } f \leq 0.4 \text{ or } f > 0.8 \\ 5f - 2 & \text{if } 0.4 < f \leq 0.6 \\ 1 & \text{if } 0.6 < f \leq 0.7 \\ -10f + 8 & \text{if } 0.7 < f \leq 0.8 \end{cases} \tag{68}$$

$$S_2(f) = \begin{cases} 0 & \text{if } f \leq 0.2 \text{ or } f > 0.6 \\ 10f - 2 & \text{if } 0.2 < f \leq 0.3 \\ 1 & \text{if } 0.3 < f \leq 0.4 \\ -5f + 3 & \text{if } 0.4 < f \leq 0.6 \end{cases} \tag{69}$$

$$S_3(f) = \begin{cases} 1 & \text{if } f \leq 0.1 \\ -5f + 1.5 & \text{if } 0.1 < f \leq 0.3 \\ 0 & \text{if } f > 0.3 \end{cases} \tag{70}$$

$$S_4(f) = \begin{cases} 0 & \text{if } f \leq 0.7 \\ 5f - 3.5 & \text{if } 0.7 < f \leq 0.9 \\ 1 & \text{if } f > 0.9 \end{cases} \tag{71}$$

and construct the membership value set $(S_1(f), S_2(f), S_3(f), S_4(f))$. The strategy selection at each iteration follows a fuzzy logic rule-based system. Let $\mathcal{A}(f) = \{i \in \{1, 2, 3, 4\} : S_i(f) \neq 0\}$ denote the set of active strategies (those with non-zero membership values) at evolutionary state $f$. The strategy $S(t)$ at iteration $t$ is selected according to:

$$S(t) = \begin{cases} \arg\max_{i \in \{1,2,3,4\}} S_i(f) & \text{if } |\mathcal{A}(f)| = 1 \\ \min\{j \in \mathcal{A}(f) : j > S(t-1)\} & \text{if } |\mathcal{A}(f)| = 2 \text{ and exists} \\ \min\{j \in \mathcal{A}(f) : j < S(t-1)\} & \text{if } |\mathcal{A}(f)| = 2 \text{ otherwise} \end{cases} \tag{72}$$

**Algorithm 1:** Metric Adaptive Particle Swarm Optimization (MAPSO)

**Input:** Loss function $\mathcal{L}(\theta)$, particles $N_p$, epochs $N_{\text{epochs}}$, dimension $D$
**Output:** Optimal parameter set $\theta^*$
`// Initialization`
**for** $\mu = 1$ **to** $N_p$ **do**
- $\theta_{\mu,i}(0) \sim \mathcal{N}(0, \sigma^2)$, $v_{\mu,i}(0) \sim U(v_{\min}, v_{\max})$ for $i = 1, \dots, D$;
- $\theta_\mu^{(\text{cog})}(0) \leftarrow \theta_\mu(0)$;

$A_{\mu\nu}(0) \leftarrow 1$ for all $\mu \neq \nu$, $\theta_\mu^{(\text{soc})}(0) \leftarrow \arg\min_\nu \mathcal{L}(\theta_\nu(0))$ for all $\mu$;
$w(0) \leftarrow 0.9$, $c_{\text{cog}}(0) \leftarrow 2$, $c_{\text{soc}}(0) \leftarrow 2$, $S(0) \leftarrow 1$;

**for** $t = 0$ **to** $N_{\text{epochs}} - 1$ **do**
- `// Swarming dynamics`
- **for** $\mu = 1$ **to** $N_p$, $i = 1$ **to** $D$ **do**
  - $v_{\mu,i}(t+1) \leftarrow w(t) v_{\mu,i}(t) + c_{\text{cog}}(t)[\theta_{\mu,i}^{(\text{cog})}(t) - \theta_{\mu,i}(t)] + c_{\text{soc}}(t)[\theta_{\mu,i}^{(\text{soc})}(t) - \theta_{\mu,i}(t)]$;
  - $\theta_{\mu,i}(t+1) \leftarrow \theta_{\mu,i}(t) + v_{\mu,i}(t+1)$;
- `// Update personal and local bests`
- **for** $\mu = 1$ **to** $N_p$ **do**
  - **if** $\mathcal{L}(\theta_\mu(t+1)) < \mathcal{L}(\theta_\mu^{(\text{cog})}(t))$ **then**
    - $\theta_\mu^{(\text{cog})}(t+1) \leftarrow \theta_\mu(t+1)$;
  - $\nu^* \leftarrow \arg\min_{\nu : A_{\mu\nu} \neq 0} \mathcal{L}(\theta_\nu^{(\text{cog})}(t+1))$;
  - **if** $\mathcal{L}(\theta_{\nu^*}^{(\text{cog})}(t+1)) < \mathcal{L}(\theta_\mu^{(\text{soc})}(t))$ **then**
    - $\theta_\mu^{(\text{soc})}(t+1) \leftarrow \theta_{\nu^*}^{(\text{cog})}(t+1)$;
- `// Update metric interaction network`
- Compute $n_{\text{neigh}}(t)$ using quartic schedule (global→local→global) and distance matrix $d_{\mu\nu}$;
- $A_{\mu\nu}(t) \leftarrow \mathbb{1}(d_{\mu\nu}(t) < d_\mu^{(n_{\text{neigh}})}(t))$ where $d_\mu^{(n_{\text{neigh}})}$ is $n_{\text{neigh}}$-th smallest distance;
- `// Compute evolutionary state and select strategy`
- Compute evolutionary state $f(t)$;
- Compute membership values $S_1(f), S_2(f), S_3(f), S_4(f)$ and select $S(t+1)$ via fuzzy logic;
- `// Update adaptive parameters based on strategy` $S(t+1)$
- Update $c_{\text{cog}}(t+1), c_{\text{soc}}(t+1)$ according to strategy $S(t+1) \in \{1, 2, 3, 4\}$;
- Clip: $c_{\text{cog}}, c_{\text{soc}} \in [3/2, 5/2]$ and normalize if sum $> 4$;
- $w(t+1) \leftarrow 1/(1 + (3/2)\exp(-13 f(t)/5))$;
- `// Mutation step (if in convergence state` $S(t+1) = 3$`)`
- **if** $S(t+1) = 3$ **then**
  - $\sigma(t+1) \leftarrow \sigma_{\max} - (\sigma_{\max} - \sigma_{\min})(t+1)/N_{\text{epochs}}$;
  - **for** $\mu = 1$ **to** $N_p$ **do**
    - Mutate random component of $\theta_\mu^{(\text{soc})}(t+1)$ with strength $\sigma(t+1)$;
    - Accept mutation if loss improves;

**return** $\arg\min_\mu \mathcal{L}(\theta_\mu^{(\text{soc})}(N_{\text{epochs}}))$

where $S(t-1)$ is the previously selected strategy, and the circular ordering wraps around (i.e., after strategy 4 comes strategy 1).

Then, the parameters are updated according to a small positive number $\delta(t) = 0.05(1 + r(t))$, with $r(t)$ a random number between 0 and 1 extracted at each step. If $f(t)$ is relatively large, the strategy should promote "exploration" at the next epoch, i.e., particles should move towards their personal best rather than the local best. Thus,

$$S(t+1) = 1 \implies c_{\text{cog}}(t+1) = c_{\text{cog}}(t) + \delta, \quad c_{\text{soc}}(t+1) = c_{\text{soc}}(t) - \delta \ . \tag{73}$$

If $f(t)$ is relatively small, the strategy should promote "exploitation", i.e., particles should slightly prefer their personal best:

$$S(t+1) = 2 \implies c_{\text{cog}}(t+1) = c_{\text{cog}}(t) + \frac{\delta}{2}, \quad c_{\text{soc}}(t+1) = c_{\text{soc}}(t) - \frac{\delta}{2} \ . \tag{74}$$

If $f(t)$ is very small, it indicates that particles are close together and the swarm is converging. The strategy should promote "convergence", slightly increasing both the cognitive and the social coefficients:

$$S(t+1)=3 \implies c_{\mathrm{cog}}(t+1)=c_{\mathrm{cog}}(t)+\frac{\delta}{2}, \quad c_{\mathrm{soc}}(t+1)=c_{\mathrm{soc}}(t)+\frac{\delta}{2}\ . \tag{75}$$

Finally, if $f$ is extremely large, it means that the swarm has split, and a "jumping-out" strategy should be promoted by increasing the social coefficient:

$$S(t+1)=4 \implies c_{\mathrm{cog}}(t+1)=c_{\mathrm{cog}}(t)-\delta, \quad c_{\mathrm{soc}}(t+1)=c_{\mathrm{soc}}(t)+\delta\ . \tag{76}$$

To avoid too large or too small coefficients, we also apply two clipping strategies, so that neither coefficient can be larger than 5/2 or smaller than 3/2. Then, if $c_{\mathrm{cog}}(t+1)+c_{\mathrm{soc}}(t+1)>4$, we renormalize them as

$$c_{\mathrm{cog}}(t+1)=4\frac{c_{\mathrm{cog}}(t+1)}{c_{\mathrm{cog}}(t+1)+c_{\mathrm{soc}}(t+1)}, \quad c_{\mathrm{soc}}(t+1)=4\frac{c_{\mathrm{soc}}(t+1)}{c_{\mathrm{cog}}(t+1)+c_{\mathrm{soc}}(t+1)}\ . \tag{77}$$

Besides this strategy-based updates of the social and cognitive coefficients, the inertia parameter is also updated according to $f(t)$:

$$w(t+1)=\frac{1}{1+3/2\exp\left(-13f(t)/5\right)}\in[0.4,0.9] \tag{78}$$

so that the inertia decreases when $f$ does.

**Local best mutations**

In the convergence state, where $f(t)$ was small at the previous epochs and the current strategy is $S(t+1)=3$, we hybridize the particle swarm dynamics with a mutation step. In particular, we attempt to mutate each local best $\theta_{\mu}^{(\mathrm{soc})}(t+1)$ along a random direction as

$$\theta_{\mu,i^*}^{(\mathrm{soc,mut})}(t+1)=\theta_{\mu,i^*}^{(\mathrm{soc})}(t+1)+r(t+1)\sigma(t+1) \tag{79}$$

where $i^*$ is a random component between 1 and $D$, $r(t+1)\sim\mathcal{N}(0,1)$, and

$$\sigma(t)=\sigma_{\max}-(\sigma_{\max}-\sigma_{\min})\frac{t}{N_{\mathrm{epochs}}} \tag{80}$$

is the mutation strength. We typically take $\sigma_{\max}=1$ and $\sigma_{\min}=0.1$, to initially favor large mutations and then reduce their strength. For each particle $\mu$, the local best is updated if and only if

$$\mathcal{L}\left(\theta_{\mu}^{(\mathrm{soc,mut})}(t+1)\right)<\mathcal{L}\left(\theta_{\mu}^{(\mathrm{soc})}(t+1)\right) \tag{81}$$

and otherwise it is discarded.

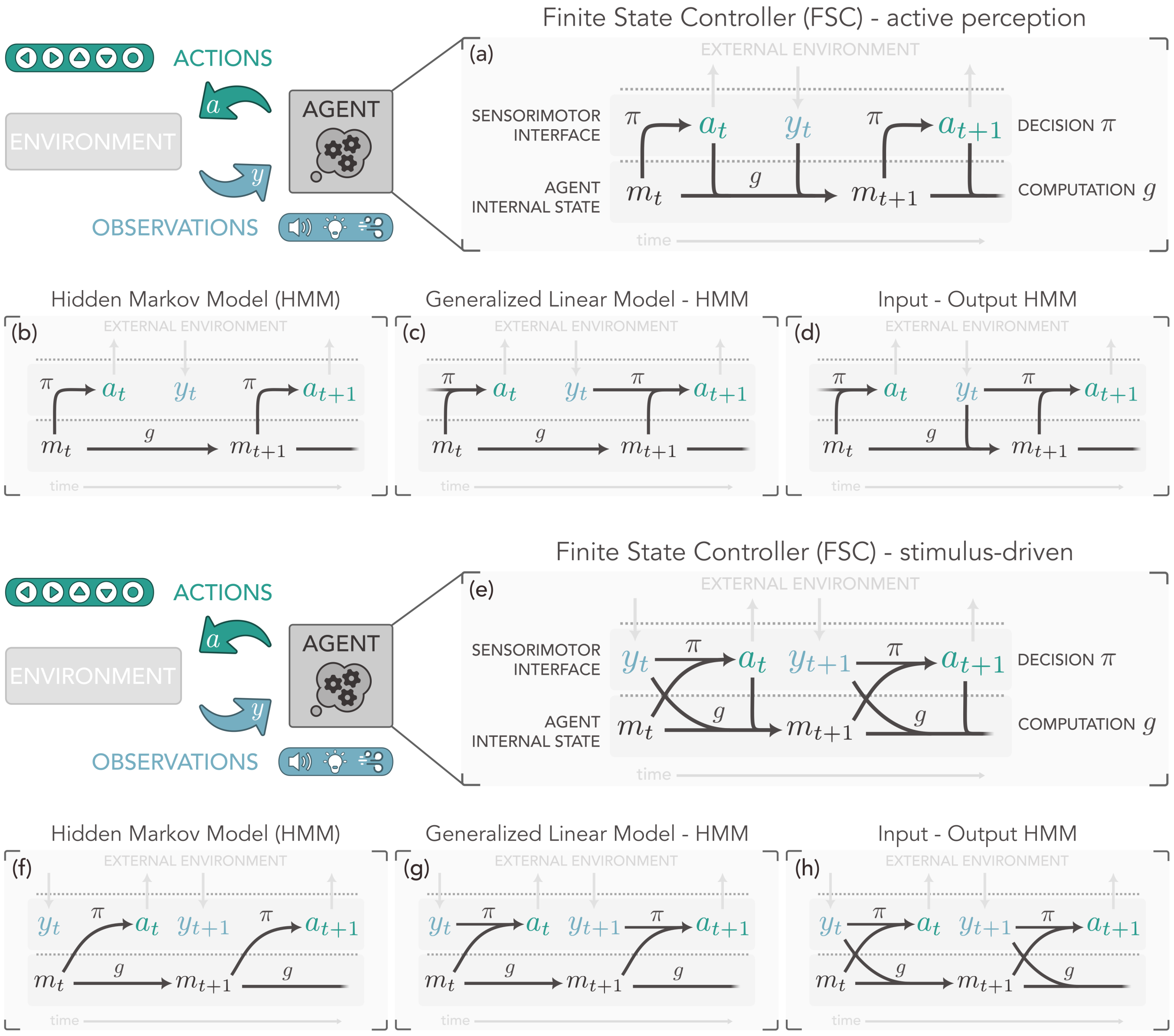


FIG. S1. Comparison between finite state controllers and other state space models in the case where observations causally follow actions (active perception) and in the case where observations are passively received from the environment and inform actions (stimulus-driven). (a) The FSC model of the agent in the case of active perception, where transitions between internal states are fully recurrent - i.e., they depend on the previous observation, action, and state. (b) In a hidden Markov model (HMM), observations are ignored, and transitions between internal states are autonomous. The policy is known as the emission probability, and solely depends on the current internal state (as in FSCs). (c) In a generalized linear model HMM (GLM-HMM), the transitions between internal states remain autonomous, but the policy depends explicitly on the observation via a generalized linear model [5]. (c) In a general input-output HMM [6], both internal transitions and the policy depend on the observation, but not on the emissions, as in FSCs. (e-h) Same as the previous panel, but for an agent that is stimulus-driven.

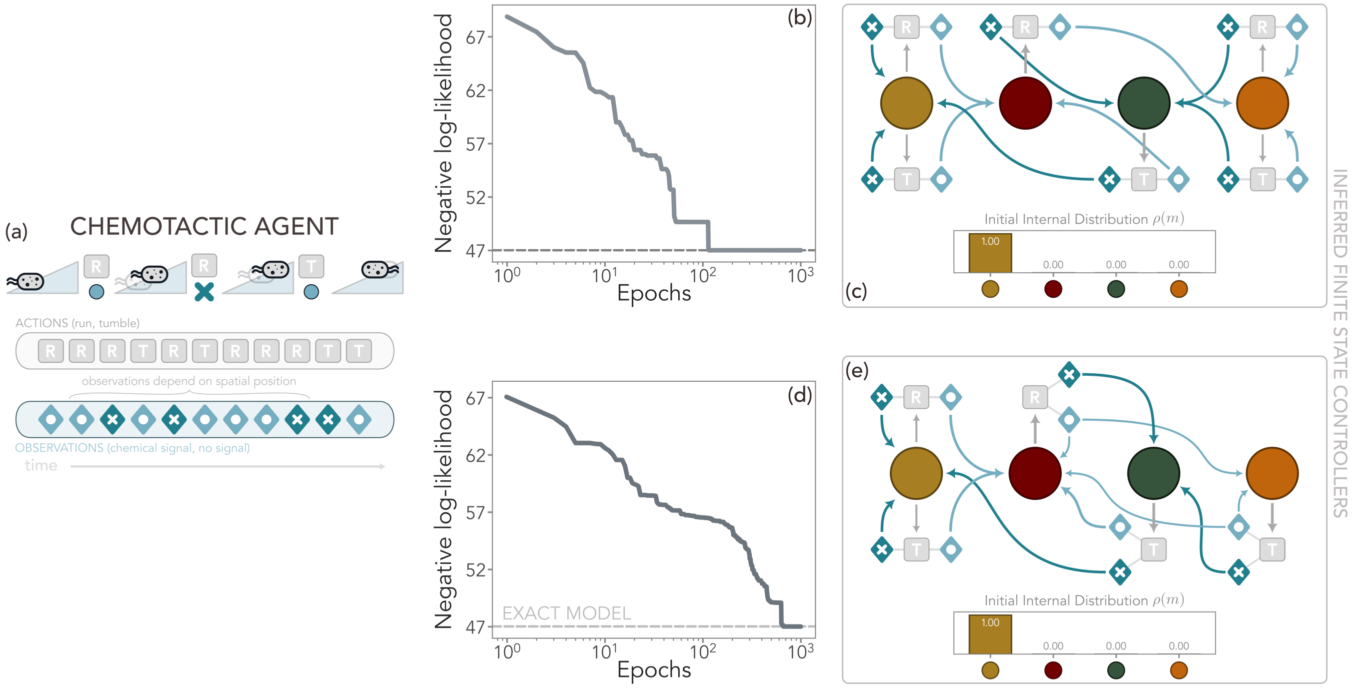


FIG. S2. Behaviorally-equivalent FSCs inferred for the chemotactic agent introduced in the main text. (a) A chemotactic agent in a $1D$ concentration gradient. At each time, the agent may sense a chemical (observation $\bullet$) or not ($\times$) with a probability that depends on its position. If in the last two steps it received the same observation, the action is chosen at random between tumble (reorienting) and run (move by a fixed amount in the current direction). Instead, if the last two observations have been $\times$ and $\bullet$, respectively, the agent runs, and otherwise it tumbles. (b-c) Negative log-likelihood during training and inferred FSC with $M = 4$ reported in the main text. In this case, the internal states of the FSC represent the four combinations of past observations used by the agent to make decisions: ● $\sim (y_{t-2}, y_{t-1}) = (\times, \times)$; ● $\sim (\times, \bullet)$; ● $\sim (\bullet, \times)$; ● $\sim (\bullet, \bullet)$. (d-e) A behaviorally-equivalent but different inferred FSC, which reaches the same negative log-likelihood as the previous one and matches that of the original agent. In this case, the stochasticity of the policy when $(y_{t-2}, y_{t-1}) = (\bullet, \bullet)$ is reproduced by equally probable internal transition between the ● and ● states, which perfectly reproduce the original diffusive behavior. This equivalence is possible because the internal computation matrices are low-rank.

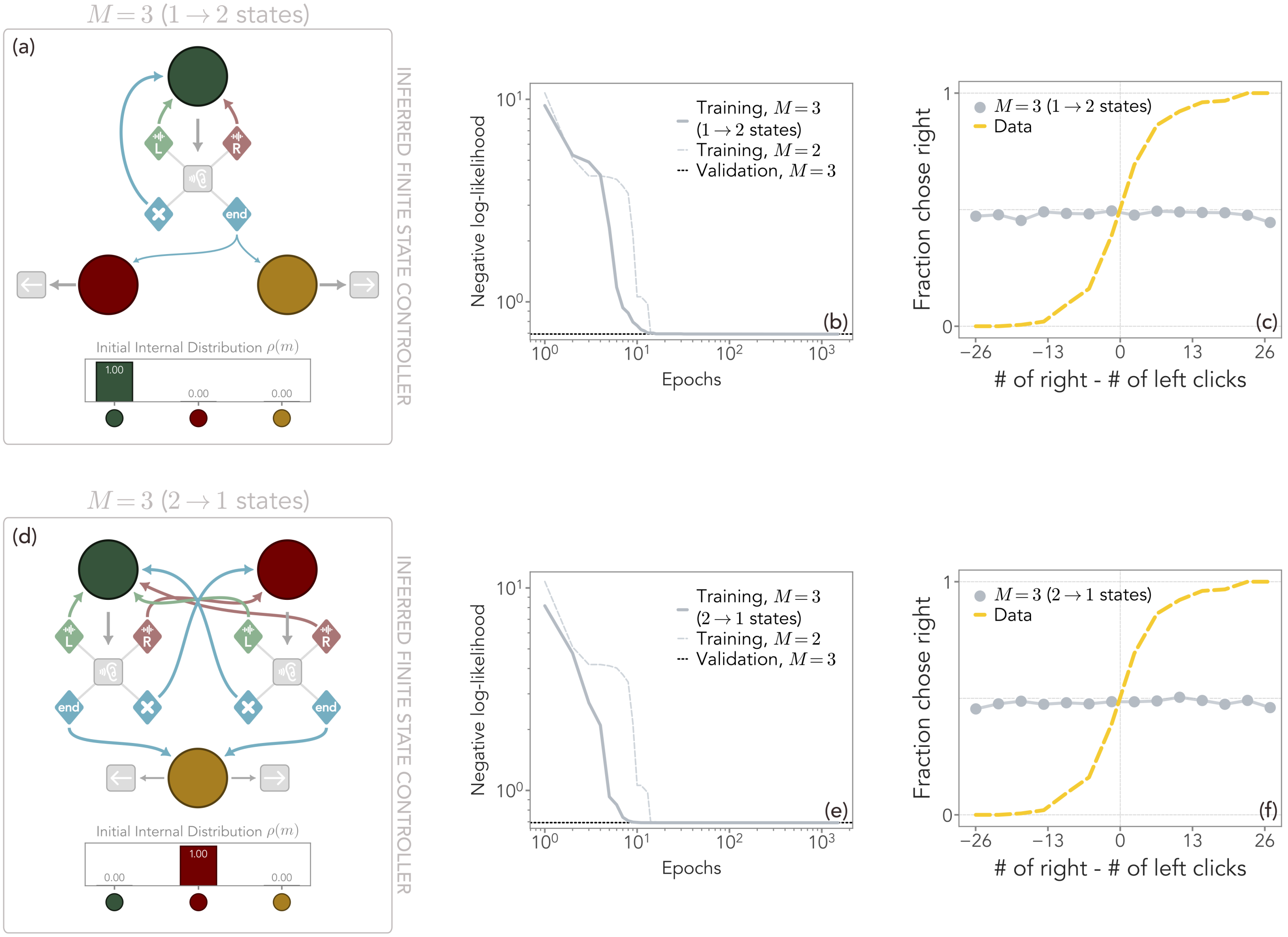


FIG. S3. Properties of the inferred FSCs with $M = 3$ for rat decision-making during evidence accumulation tasks (see Figure 3 of the main text). Two classes of equivalent FSCs with $M = 3$ exist, and suggest that the behavior can be recapitulated with a two-layer structure for the internal states and for $M \geq 4$. As in the main text, states in the first layer take the "listen" action only, whereas those in the second layer take the "go" actions. (a-c) A first possibility is a two-layer structure with one single node in the first layer ( ) and two in the second ( and ). The two states in the second layer specialize in taking the two final actions, i.e., going either right or left, after the end observation is received and the trial terminates. In this way, however, no computation can be performed in the remaining state . Thus, in terms of behavior, this topology is fully equivalent to the $M = 2$ one, and indeed it achieves the same negative log-likelihood on the training and validation set (b), as well as acting randomly regardless of the number of left and right observations it received (c). (d-e) The second option is an FSC with two states in the first layer ( and ) and one in the second ( ). In principle, the nodes in the first layer can now perform computations, as in the $M = 4$ case. However, regardless of the computation, when the end observation is received, the state must switch to and is once again forced to act randomly. In this Figure, no a priori structure was imposed for inference.

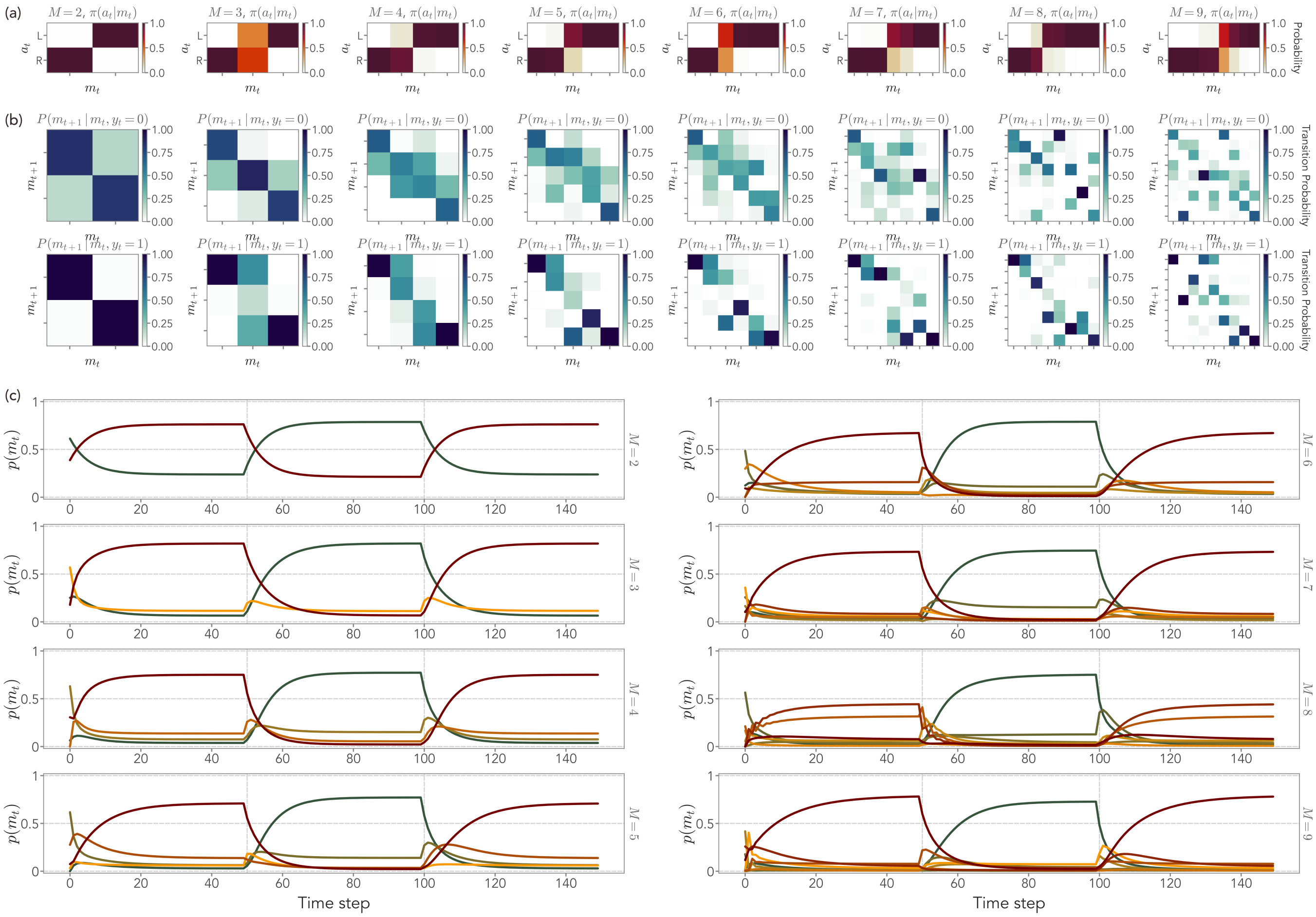

FIG. S4. Properties of the inferred FSCs for mice decision-making in changing environments for different values of $M$, from $M = 2$ to $M = 9$. From $M = 4$ onward, the negative log-likelihood reaches a plateau (see Figure 5 of the main text). (a) Inferred policies. In all cases, the internal states roughly partition into two groups, each of them taking one of the two available actions (go left or go right). (b) Probability of transitioning from one internal state to the other, if the reward is given ($y = 0$) or not ($y = 1$). As in the main text, a linear structure is clearly visible up to $M = 4$. For $M \geq 5$, the FSC topology remains linear, although several internal states may not be used as the behavior is accurately reconstructed with fewer nodes. (c) Probability of occupying a given internal state for a fixed sequence of ports (see Figure 5g-i of the main text). For $M \geq 3$, switch detectors emerge. For large $M$, the same internal states may perform the same function – e.g., for $M = 8$ two internal states encode the mice's decisions when the high port has been on the right. As the likelihood suggests, this leads to an equivalent behavior to the case of $M = 4$, which already contains the minimal number of internal states to recapitulate the mice's decision process: two asymmetric switch detectors, and two states for the two ports.

[1] B. Friedland, *Control system design: an introduction to state-space methods* (Courier Corporation, 2012).
[2] C. E. Schroeder, D. A. Wilson, T. Radman, H. Scharfman, and P. Lakatos, Dynamics of active sensing and perceptual selection, Current Opinion in Neurobiology **20**, 172 (2010).
[3] J. Gottlieb, P.-Y. Oudeyer, M. Lopes, and A. Baranes, Information-seeking, curiosity, and attention: computational and neural mechanisms, Trends in Cognitive Sciences **17**, 585 (2013).
[4] R. S. Sutton, A. G. Barto, *et al.*, *Reinforcement learning: An introduction*, Vol. 1 (MIT press Cambridge, 1998).
[5] Z. C. Ashwood, N. A. Roy, I. R. Stone, I. B. Laboratory, A. E. Urai, A. K. Churchland, A. Pouget, and J. W. Pillow, Mice alternate between discrete strategies during perceptual decision-making, Nature Neuroscience **25**, 201 (2022).
[6] Y. Bengio and P. Frasconi, Input-output hmms for sequence processing, IEEE Transactions on Neural Networks **7**, 1231 (1996).
[7] J. E. Hopcroft, R. Motwani, and J. D. Ullman, *Introduction to Automata Theory, Languages, and Computation*, 3rd ed. (Addison Wesley, 2006).
[8] F. van Ede, A. G. Board, and A. C. Nobre, Goal-directed and stimulus-driven selection of internal representations, Proceedings of the National Academy of Sciences **117**, 24590 (2020).
[9] K. V. B. Verano, E. Panizon, and A. Celani, Olfactory search with finite-state controllers, Proceedings of the National Academy of Sciences **120**, e2304230120 (2023).
[10] S. C.-H. Yang, D. M. Wolpert, and M. Lengyel, Theoretical perspectives on active sensing, Current Opinion in Behavioral Sciences **11**, 100 (2016).
[11] H. Ito, S.-I. Amari, and K. Kobayashi, Identifiability of hidden markov information sources and their minimum degrees of freedom, IEEE Transactions on Information Theory **38**, 324 (1992).
[12] J. Chung, C. Gulcehre, K. Cho, and Y. Bengio, Empirical evaluation of gated recurrent neural networks on sequence modeling, arXiv preprint arXiv:1412.3555 (2014).
[13] M. Ravanelli, P. Brakel, M. Omologo, and Y. Bengio, Light gated recurrent units for speech recognition, IEEE Transactions on Emerging Topics in Computational Intelligence **2**, 92 (2018).
[14] M. Usher and J. L. McClelland, The time course of perceptual choice: the leaky, competing accumulator model., Psychological Review **108**, 550 (2001).
[15] Z.-H. Zhan, J. Zhang, Y. Li, and H. S.-H. Chung, Adaptive particle swarm optimization, IEEE Transactions on Systems, Man, and Cybernetics, Part B (Cybernetics) **39**, 1362 (2009).
